# Visualizing protein-protein interactions in plants by rapamycin-dependent delocalization

**DOI:** 10.1101/2020.03.09.983270

**Authors:** Joanna Winkler, Evelien Mylle, Andreas De Meyer, Benjamin Pavie, Julie Merchie, Peter Grones, Daniël Van Damme

## Abstract

Identifying protein-protein interactions (PPI) is crucial for understanding biological processes. Many PPI tools are available, yet only some function within the context of a plant cell. Narrowing down even further, only a few tools allow complex multi-protein interactions to be visualized. Here, we present a conditional *in vivo* PPI tool for plant research that meets these criteria. Knocksideways in plants (KSP) is based on the ability of rapamycin to alter the localization of a bait protein and its interactors via the heterodimerization of FKBP and FRB domains. KSP is inherently free from many limitations of other PPI systems. This *in vivo* tool does not require spatial proximity of the bait and prey fluorophores and it is compatible with a broad range of fluorophores. KSP is also a conditional tool and therefore the visualization of the proteins in the absence of rapamycin acts as an internal control. We used KSP to confirm previously identified interactions in *Nicotiana benthamiana* leaf epidermal cells. Furthermore, the scripts that we generated allow the interactions to be quantified at high throughput. Finally, we demonstrate that KSP can easily be used to visualize complex multi-protein interactions. KSP is therefore a versatile tool with unique characteristics and applications that complements other plant PPI methods.

## INTRODUCTION

Unraveling macromolecular complexes and interaction networks among proteins plays a pivotal role in understanding the complexity of biological processes. In the past 20 years, the number of articles that feature protein-protein interactions (PPIs) as a topic or focusing on PPI technology development has been steadily increasing (Xing et al., 2016). Although a plethora of tools to investigate PPIs are currently available (Miura, 2018; Titeca et al., 2019; Wiens and Campbell, 2018), relatively few can be used *in planta* (Fukao, 2012; Lampugnani et al., 2018; Struk et al., 2019; Xing et al., 2016). In plant systems, including *Arabidopsis thaliana*, multiple interactomes have been described based on large-scale screenings (Arabidopsis Interactome Mapping Consortium, 2011; Boruc et al., 2010a; Jones et al., 2014; Piya et al., 2014). Interactomes plot the landscape surrounding a certain bait protein, yet seldom provide information on specific or direct interaction partners. Evaluating specific biological functions and the mode of action of a protein of interest usually requires multi-technical approaches (Struk et al., 2019). Thus, selecting the optimal PPI tools to use represents a crucial task.

Affinity-purification (AP), including co-immunoprecipitation (Co-IP), tandem affinity purification and proximity labelling, followed by mass spectrometry (MS) experiments (Arora et al., 2020; Masters, 2004; Ransone, 1995; Rigaut et al., 1999) are powerful screening tools to identify PPIs. One advantage of these tools is that the interactions occur in their physiological environment (Rubio et al., 2005; Xing et al., 2016). However, AP/MS requires a cell lysis step, during which it is possible to impose false-positive PPIs, and even more common, to disrupt weak ones (Miteva et al., 2013). On the contrary, with proximity labelling methods, there is no need to preserve PPIs during lysis and extraction. Proximity-dependent biotin identification (BioID) allows the spatial conditions of PPIs to be preserved and is particularly useful for identifying low-affinity and transient interactions (Roux et al., 2012), including in plants (Arora et al., 2020; Branon et al., 2018; Conlan et al., 2018; Das et al., 2019; Khan et al., 2018; Lin et al., 2017; Mair et al., 2019; Zhang et al., 2019). The downside of the above techniques is that they do not provide any information about the binary interactions occurring between bait and prey proteins.

Binary interaction assays using the yeast *Saccharomyces cerevisiae*, such as the yeast two-hybrid (Y2H) system (Fields and Song, 1989; Wen, 2014), rely on the reconstitution of bipartite transcriptional activators (Lampugnani et al., 2018). Y2H can be upgraded into Yeast three-hybrid (Y3H), which brings the system to the tripartite-protein interaction level (Alberti et al., 2007; Cottier et al., 2011; Maruta et al., 2016). Also, incorporating Y2H with next-generation sequencing or recombination based Y2H ‘library vs. library’ screening (Erffelinck et al., 2018; Yang et al., 2018) resulted in the development elaborate, high-throughput PPI screening tools. Nevertheless, as yeasts lack certain chaperones or posttranslational modifications that occur *in planta*, identifying interactions, which rely on these, is not feasible. In addition, proteins might require several additional proteins to stabilize their interaction. For instance, the octameric endocytic TPLATE complex was shown to be a very robust multi-subunit complex via proteomics analysis, but only one interaction was identified in yeast (Gadeyne et al., 2014). Therefore, because yeast assays rely on a heterologous system lacking expression of other plant proteins beside the bait and prey, they have their limitations. These assays also do not provide information about the subcellular localization of the PPI.

Two commonly used *in planta* interaction techniques that confer subcellular localization information are Bimolecular Fluorescence Complementation (BiFC) and Förster resonance energy transfer (FRET). BiFC is a versatile tool that relies on the ability of complementary halves of fluorescent proteins (FP) to reconstitute their properties upon PPI (Ghosh et al., 2000). BiFC has been adapted for plant systems (Bracha - Drori et al., 2004; Hu et al., 2002; Walter et al., 2004), optimized for ratiometric imaging (Grefen and Blatt, 2012), and developed into tripartite split-GFP in order to avoid self-assembly (Cabantous et al., 2013) or to visualize triple interactions (Offenborn et al., 2015). Weak and transient PPIs can be visualized via these methods, as once FP halves re-assemble, the interaction is locked. The interaction is thus irreversible, and interacting proteins are artificially stabilized and remain in close proximity (Magliery et al., 2005). While this is beneficial for identifying weak interactions, there are also several major limitations to this system. The stabilization of the interaction, regardless of the localization and the time of interaction, might prevent linking of the localization to the site of interaction. This technique also requires testing multiple combinations of orientations of the split-GFP proteins tags, and it is difficult to use the appropriate negative controls (Kudla and Bock, 2016).

FRET is one of the *in vivo* assays *in planta* where interplay between two proteins of interest can be measured via, among others, fluorescence lifetime imaging (FLIM) (Förster, 1948; Gadella et al., 1993; Sun et al., 2011; Xing et al., 2016). FRET-FLIM has recently been developed into a three-fluorophore system that allows ternary PPIs to be visualized (Glöckner et al., 2019). Nevertheless, FRET-FLIM is quite labor-intensive and requires careful interpretation and multiple control assays. Moreover, is it significantly limited by the intrinsic properties of the FPs (Bhat et al., 2006; Struk et al., 2019).

Here, we present a PPI tool we named Knocksideways in plants (KSP), which can be compared to an intracellular Co-IP experiment. This tool relies on the rapamycin-induced heterodimerization between the FKBP domain of HsFKBP12 (FKBP) and the FKBP12 rapamycin-binding domain of mTOR (FRB). This dimerization tool is well-established and has successfully been used in human, animal, plasmodium, and yeast systems (Haruki et al., 2008; Putyrski and Schultz, 2012; Wood et al., 2017; Chojnowski et al., 2018; Hughes and Waters, 2017; Nomura et al., 2018; Putyrski and Schultz, 2012) and has occasionally been used in plants (Li et al., 2011; Rosenfeldt et al., 2008). We were inspired by the Knocksideways (KS) approach in animal cells, which combines conditional FKBP-FRB binding with rerouting proteins to the mitochondria (Robinson et al., 2010). This type of ‘drag-away’ technique has already been explored for PPIs in plant research. For instance, the Nuclear Translocation Assay (NTA), the Cytoskeleton-based localization Assay for Protein-Protein Interaction (CAPPI), and the Modification of Intracellular Localization (MILo) function via a similar approach (Dixon and Lim, 2010; Kaplan - Levy et al., 2014; Lv et al., 2017). Nevertheless, the above tools are not conditional, and their subcellular targeting options reported so far are limited. In this study, we focused on introducing KS *in planta*, expanding the availability of the subcellular FRB anchors, and developing the system into a PPI tool for examining complex multiprotein interactions where the results can be quantified using *Nicotiana benthamiana* leaf epidermal cells.

## RESULTS

### Rapamycin efficiently relocalizes FKBP-fusions to FRB-tagged subcellular compartments in planta

First, we tested if rapamycin-induced FKBP-FRB* domain heterodimerization is efficient in plants. To do this, we amplified the FRB* domain (∼84 kDa, together with TagBFP2) from MAPPER (Chang et al., 2013) and fused it with various peptides, anchoring it at different subcellular locations. This enriches the tool’s application for diversely localized proteins of interest. *Arabidopsis thaliana* MICROTUBULE END BINDING PROTEIN EB1A (AtEB1a), a plus-end microtubule marker (Van Damme et al., 2004), was used to design the microtubular anchor EB1a-TagBFP2-FRB*. We fused the FRB* domain to a nuclear localization signal (NLS) to create the nuclear-localized anchor NLS-TagBFP2-FRB*. The Nterminal part of the Ca^2+^-dependent protein kinase CPK34, containing a myristoylation/palmitoylation signal responsible for plasma membrane (PM) localization (Kirik et al., 2012; Myers et al., 2009), was used to create the MYRI-TagBFP2-FRB* PM anchor. Finally, for mitochondrial targeting, we used the transit sequence of the yeast mitochondrial outer membrane protein Tom70p (MITO-TagBFP2-FRB*), previously reported to be functional in mammalian cells (Kessels and Qualmann, 2002; Robinson et al., 2010). All newly generated constructs are available as multisite-Gateway building blocks (Supplemental Figure 1).

We expressed the aforementioned FRB* constructs in *Nicotiana benthamiana* abaxial leaf cells and used spinning disk confocal microscopy to visualize the fusion proteins. All FRB* constructs were present at their expected subcellular localizations. We then combined these constructs with assorted mCherry-FKBP-tagged bait proteins (bait + ∼86 kDa) and monitored their localization prior to and after rapamycin treatment.

TPLATE is a subunit of the octameric TPLATE complex (TPC) involved in clathrin-mediated endocytosis (Gadeyne et al., 2014). In *N. benthamiana*, over-expressed TPLATE localizes ubiquitously in the cytoplasm (Gadeyne et al., 2014), which is similar to the results for our TPLATE-mCherry-FKBP. After rapamycin treatment, TPLATE was rerouted to the microtubules (MTs) in the presence of EB1a-TagBFP2-FRB* (Figure 1A). MICROTUBULE-ASSOCIATED PROTEIN 65-1 (MAP65-1) is a MT cross-linker (Tulin et al., 2012; Van Damme et al., 2004). In the presence of the microtubule depolymerizing agent oryzalin, rapamycin can delocalize and trap MAP651 in the nucleus via binding to NLS-TagBFP2-FRB* (Figure 1B). Similar to what has been reported before (Boruc et al., 2010a; Spinner et al., 2013), mCherry-FKBP fusions of the protein phosphatase 2A regulatory subunit RCN1 and the cyclin-dependent kinase CELL DIVISION CONTROL 2 (CDKA;1) were largely cytoplasmic in the absence of the dimerizer. Rapamycin delocalized RCN1 to the plasma membrane-docked MYRITagBFP2-FRB* (Figure 1C) and pulled CDKA;1 to the mitochondria in the presence of MITO-TagBFP2-FRB* (Figure 1D).

**Figure 1.**
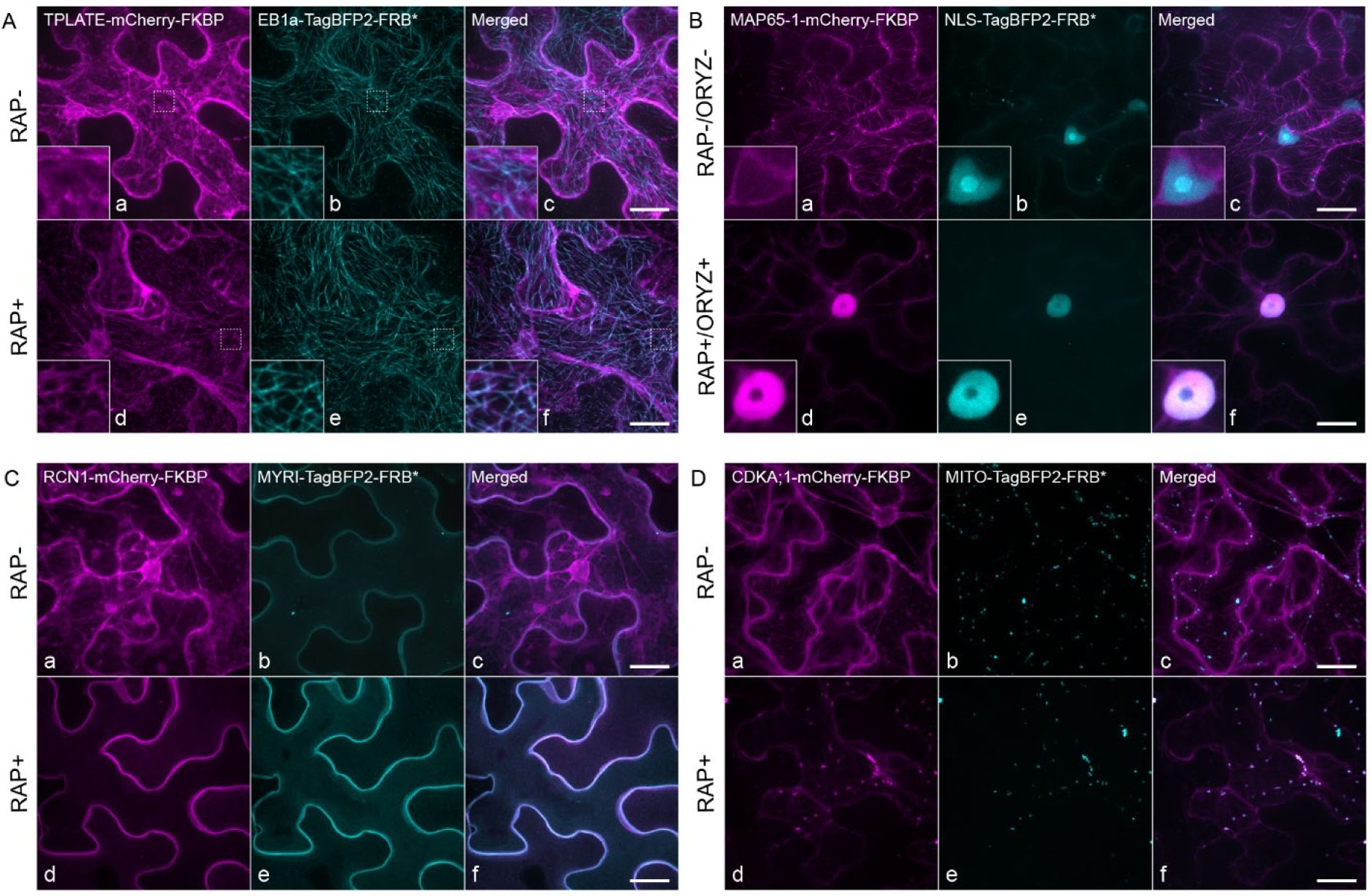
Rapamycin allows FKBP protein fusions to be relocalized to different subcellular compartments in *Nicotiana benthamiana* leaf epidermal cells. Representative Z-stack projections of epidermal *N. benthamiana* cells transiently expressing mCherry-FKBP and TagBFP2-FRB* fusion proteins in the absence (RAP-) or presence (RAP+) of rapamycin. (A) Targeting to microtubules. Without rapamycin, TPLATE-mCherry-FKBP localizes to the cytoplasm (a), while EB1a-TagBFP2-FRB* decorates microtubules (b). Rapamycin redirects TPLATE-mCherry-FKBP effectively to the microtubule cytoskeleton (d-f). The insets represent single-layer enlarged, contrast-enhanced microtubular areas. (B) Targeting to the nucleus. Without rapamycin and oryzalin (RAP-/ORYZ-), MAP65-1 labels the cortical microtubule cytoskeleton and is excluded from the nucleus (a). NLS-TagBFP2-FRB* strongly accumulates inside the nuclei (b). Oryzalin releases MAP65-1-mCherry-FKBP into the cytosol and rapamycin relocalizes it to the nuclei (RAP+/ORYZ+, d-f). The insets represent single-layer enlarged nuclear areas. (C) Targeting to the plasma membrane. Without rapamycin, RCN1-mCherry-FKBP localizes to the cytoplasm and nuclei (a), while MYRI-TagBFP2-FRB* is exclusively located at the plasma membrane (b). Rapamycin relocalizes RCN1-mCherry-FKBP effectively to the plasma membrane (d-f). (D) Targeting to mitochondria. Without rapamycin, CDKA;1-mCherry-FKBP localizes predominantly in the cytoplasm (a) and does not co-localize with the mitochondrial targeted MITO-TagBFP2-FRB* (b). Following rapamycin treatment, CDKA;1-mCherry-FKBP is redirected to the mitochondria (d-f). Images on the right contain the merged panels (c, f). Scale bars = 20μm.

We observed that MITO-TagBFP2-FRB*-labeled structures differed in size and occasionally clustered into larger structures. To confirm that the punctate MITO-TagBFP2-FRB* signals indeed represented mitochondria, we expressed MITO-TagBFP2-FRB* in *N. benthamiana* leaves and stained them with the mitochondrial dye MitoTracker™ Red. The staining colocalized with MITO-TagBFP2-FRB* regardless of the clustering (Supplemental Figure 2A). Additionally, to exclude the possibility that clustering might affect the localization of FKBP-fused protein, we co-infiltrated *N. benthamiana* leaves MITO-TagBFP2-FRB* with the mCherry-FKBP-tagged transcription factor LONESOME HIGHWAY (LHW-mCherry-FKBP). In the presence of rapamycin, LHW-mCherry-FKBP was rerouted to the mitochondria irrespective of their aggregation (Supplemental Figure 2B).

Taken together, these results indicate that heterodimerization of FKBP-FRB* upon rapamycin addition works efficiently in plants and that the system is capable of relocalizing baits to different subcellular FRB*-anchors.

### FKBP-FRB* rapamycin-dependent binding is partially reversible

Ascomycin competes with rapamycin for the HsFKBP binding site (Paulmurugan et al., 2004) and has been shown to function in Arabidopsis mesophyll protoplasts (Li et al., 2011). We therefore wanted to test if the rapamycin-dependent delocalization of the bait protein could be reversed in the *N. benthamiana* system. We co-infiltrated leaves with TPLATE-mCherry-FKBP and the nuclear-FRB* anchor and treated the leaves with rapamycin. One hour later, we treated half of the samples with ascomycin. We imaged the leaves and quantified the nucleus/cytoplasm ratio 5-7h after the treatment, and again after approximately 24 hours (Supplemental Figure 3).

A few hours of ascomycin treatment did not revert the delocalization of TPLATE into the nucleus (Supplemental Figure 3 B, C and F). However, prolonged treatment (approximately 25 hours) reverted the nucleus/cytoplasm TPLATE intensity ratio to that of the untreated samples, in contrast to the ratio in rapamycin-only treated cells (Supplemental Figure 3 D, E and F). These results suggest that KSP using *N. benthamiana* infiltration is partially reversible. However, it is not efficient and requires prolonged ascomycin treatment.

### Knocksideways allows enzymatic reactions to be compartmentalized

We wondered whether KSP-dependent targeting of FKBP-fused proteins would allow specific enzymatic reactions to be directed. The phosphoinositide composition of the membrane serves as a landmark code, and this is crucial for the selective recruitment of proteins (Doumane and Caillaud, 2020; Noack and Jaillais, 2017). Phosphatidylinositol-4-phosphate (PI4P) can be depleted from the PM by targeting the yeast SUPPRESSOR OF ACTIN 1 (Sac1) PI4P phosphatase to this location (Doumane and Caillaud, 2020). We fused the Sac1 phosphatase and its catalytically dead version, Sac1-dead (Sac1-d) to our mCherry-FKBP module and expressed them in *N. benthamiana*, alongside our myristoylation-FRB* PM anchor and a marker for PI4P (mCitrine-P4M). Rapamycin addition targeted both phosphatases to the PM. This only resulted in a partial delocalization of the PI4P marker to discrete endosomes in the cytoplasm when the active phosphatase was used (Figure 2A-C). Our results demonstrate that KSP has potential for conditional physiological applications *in planta*.

**Figure 2.**
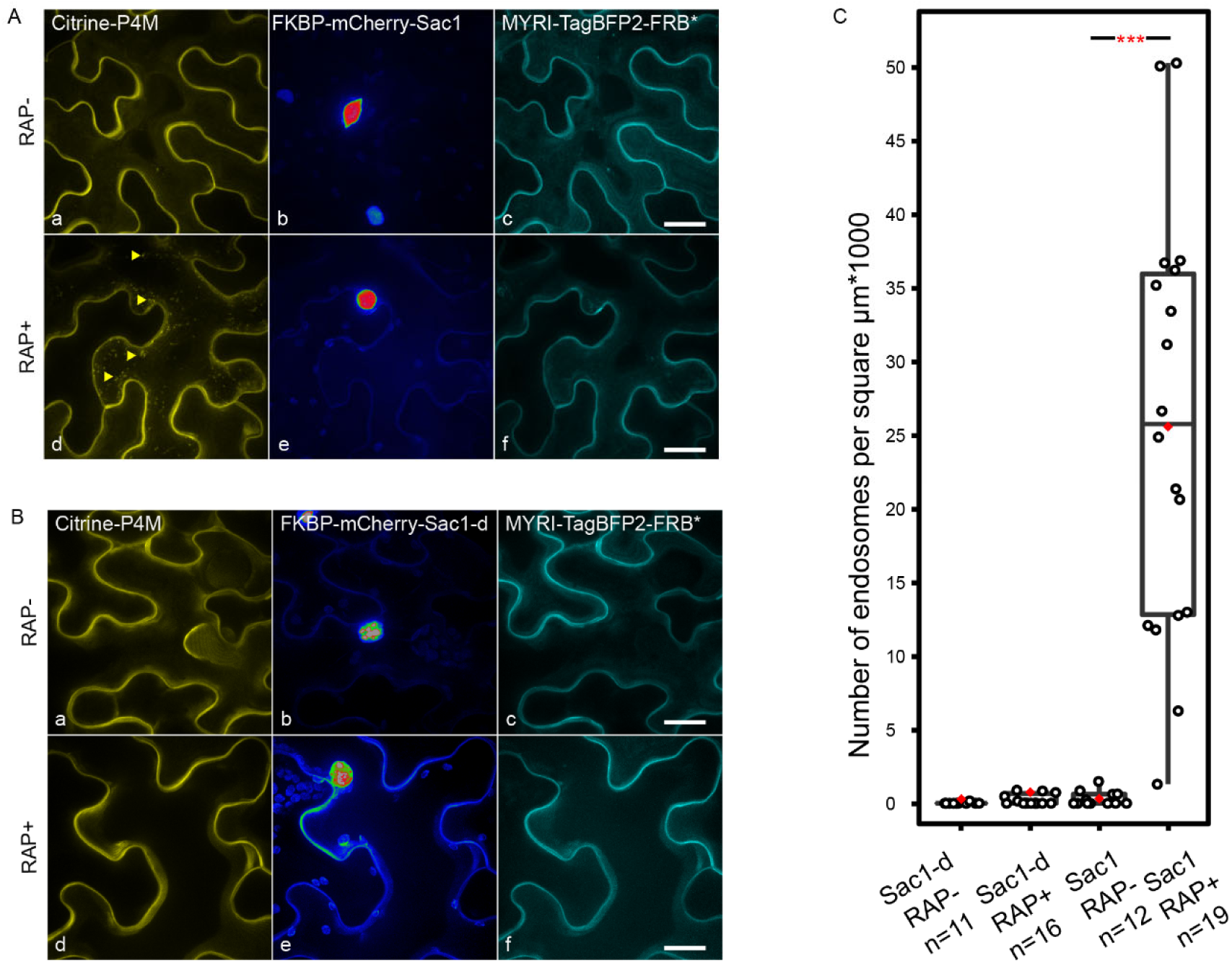
KSP allows PI4P distribution to be perturbed via the relocalization of this phosphatase to the plasma membrane. (A-B) Representative Z-stack projected images of epidermal *N. benthamiana* cells transiently expressing the PI4P sensor Citrine-P4M (a, d), the FKBP-mCherry-tagged active (A) or catalytically dead (B) yeast SUPPRESSOR OF ACTIN 1 PI4P phosphatase domain (b, e, Sac1 and Sac1-d respectively) and the PM anchor MYRI-TagBFP2-FRB* (c, f). In the absence of rapamycin (RAP-), PI4P, visualized by the Citrine-P4M sensor, localizes to the PM. Upon rapamycin treatment (RAP+), Sac1 is partially delocalized to the PM, which correlates with the relocalization of the PI4M marker at the endosomes (A panel d, yellow arrowheads). By contrast, Sac1-dead does not perturb PI4P localization (B panel d). Scale bars = 20μm. (C) Statistical analysis showing the density of endosomal delocalization upon rapamycin treatment. The black line represents the median and the red diamond represents the mean of the analyzed values. Each dot represents an individual cell, and n refers to the total number of analyzed cells. The P-value <0.001 is represented as ***. Error bars represent the 95%-confidence interval.

### Knocksideways is a robust, quantitative protein-protein interaction tool *in planta*

We then used KSP to visualize PPIs in *N. benthamiana*. To do so, we targeted our FRB* domain to a specific subcellular location. We fused several bait proteins, with different subcellular localizations and from different functional classes, to mCherry-FKBP. We expressed these fusion proteins, together with their GFP-fused interaction partners, and monitored their localization in the absence and presence of rapamycin. We designed Fiji/ImageJ-compatible scripts to easily quantify nuclear and mitochondrial delocalization upon rapamycin addition (see Methods and Supplemental File 1 for a detailed description of how to use the scripts).

To demonstrate that rapamycin itself does not cause arbitrary binding or random delocalization of proteins, we co-infiltrated leaves with free eGFP together with TPLATEmCherryFKBP and MITO-TagBFP2-FRB*. Without rapamycin, both eGFP and TPLATE-mCherry-FKBP localized ubiquitously to the cytosol. Rapamycin addition effectively delocalized TPLATE to the mitochondria and mitochondrial clusters but did not affect the localization of eGFP (Figure 3A). We quantified the mitochondria/cytoplasm intensity ratio in untreated and rapamycin-exposed samples using our MitTally script (i.e. Mitochondria Tally marks; see Supplemental File 1 and 2). Statistical analysis indicated that in contrast to eGFP, TPLATE intensity at the mitochondria was significantly increased after rapamycin treatment (Figure 3A i).

**Figure 3.**
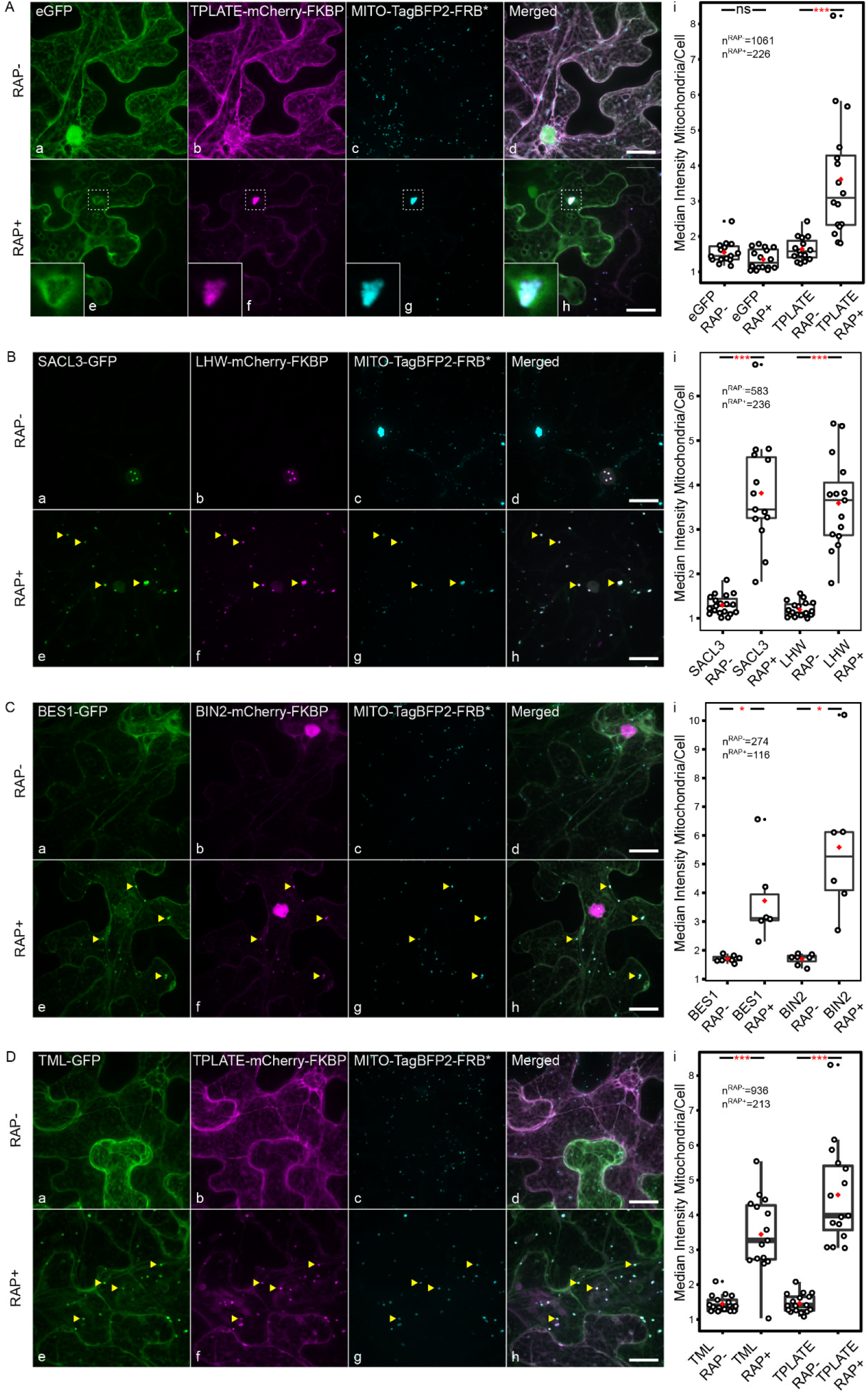
KSP allows quantitative visualization of protein-protein interactions to be performed in plants via delocalization to mitochondria. (A-D) Representative Z-stack projected images of epidermal *N. benthamiana* cells transiently expressing various GFP-fused proteins (a, e), mCherry-FKBP-fused proteins (b, f) and TagBFP2-FRB* construct targeted to the mitochondria (c, g). Panels on the right represent merged images (d, h). Free eGFP does not co-localize with the mCherry-FKBP and TagBFP2-FRB*-positive mitochondria, some of which are clustered both in the absence (RAP-) and presence (RAP+) of rapamycin (A). In the absence of rapamycin (RAP-), none of the -GFP or -mCherry-FKBP fused proteins co-localize with the TagBFP2-FRB* constructs. After rapamycin treatment, FKBP protein fusions are effectively delocalized to the mitochondria (B-D, yellow arrowheads), confirming their interaction with the FKBP-fused bait protein. The insets represent an enlarged, single-layer cytosolic area surrounding a mitochondrial cluster (A). Scale bars = 20μm. n^RAP-/RAP+^ refers to the total number of analyzed mitochondria in untreated (RAP-) or rapamycin-treated (RAP+) cells. Statistical analysis (panels i) shows the quantification of mitochondria/cytoplasm intensity ratios of co-infiltrated prey-GFP and bait-mCherry-FKBP protein fusions before and after rapamycin treatment. The black line represents the median and the red diamond represents the mean of the analyzed values. Each dot represents an individual cell. P-values <0.001 are represented as *** and <0.05 as *. Not statistically significant results are noted as ‘ns’. Error bars represent the 95%-confidence interval.

SAC51-LIKE (SACL) is a bHLH transcription factor that heterodimerizes with LONESOME HIGHWAY (LHW) (Vera-Sirera et al., 2015). The SACL/LHW interaction has been demonstrated via various conventional PPI methods, including immunoprecipitation followed by tandem mass spectrometry (IP-MS/MS), Y2H, and FRET-FLIM (De Rybel et al., 2013; Ohashi-Ito and Bergmann, 2007; Vera-Sirera et al., 2015). Here, we tested SACL-GFP and LHW-mCherry-FKBP in KSP with MITO-TagBFP2-FRB*. Without rapamycin, SACL and LHW localized to the nucleus. Rapamycin treatment delocalized both proteins to the mitochondria, thereby confirming their interaction in our system and independent of the nuclear environment of the bait and the prey protein (Figure 3B).

Phosphorylation of the transcription factor BRI1-EMS-SUPPRESSOR 1 (BES1) by the BRASSINOSTEROID-INSENSITIVE 2 (BIN2) kinase is a well-established interaction in the Brassinosteroid signaling pathway (Anwar et al., 2018; Planas-Riverola et al., 2019). We co-infiltrated *N. benthamiana* leaves with BES1-GFP with BIN2-mCherry-FKBP and MITOTagBFP2FRB*. Without rapamycin, BIN2 primarily localized to the nucleus while BES1 was ubiquitously present in the cytoplasm. Rapamycin locked BIN2 at the mitochondria where it recruited BES1 (Figure 3C).

The TPLATE complex core consists of four subunits: TPLATE, TML, LOLITA, and TASH3. Interactions between particular subunits have been demonstrated via different PPI methods (Gadeyne et al., 2014; Hirst et al., 2014; Yperman et al., 2020). We tested the TPLATE-TML interaction, as previously revealed via BiFC and Co-IP (Gadeyne et al., 2014), using KSP. Both TPLATE-mCherry-FKBP and TML-GFP were largely cytoplasmic when over-expressed in *N. benthamiana*. Exposure to rapamycin rerouted TPLATE together with TML to the mitochondria, reaffirming their interaction. The interaction was robust, and the mitochondria/cell intensity ratio was significantly altered by rapamycin treatment (Figure 3D).

To test if visualization of PPI via KSP is compatible with other subcellular locations, we investigated the efficiency of NLS-TagBFP2-FRB* as an anchor. Rapamycin rerouted TPLATE-mCherry-FKBP into the nucleus, together with TML-GFP (Figure 4A). We quantified the nucleus/cytoplasm intensity ratio in untreated and rapamycin-exposed samples using our NucTally script (i.e. Nuclear Tally marks; see Supplemental File 1 and 3). The nucleus/cytoplasm median intensity ratio, indicative of the accumulation of TPLATE/TML in the nucleus, increased significantly upon rapamycin treatment (Figure 4A i). Next, we co-infiltrated leaf cells with MAP65-1-GFP and MAP65-1-mCherry-FKBP and NLSTagBFP2-FRB*. Prior to rapamycin treatment, both MAP65-1 fusion proteins localized predominantly on microtubules, although the GFP-fusion displayed an increased propensity for aggregation compared to the mCherry-FKBP fusion, likely due to overexpression levels. After treatment with rapamycin and oryzalin, MAP651 fusion proteins were released from the microtubules and significantly co-delocalized to the nucleus (Figure 4B and Figure 4B i), while oryzalin treatment alone did not force these proteins into the nucleus (Supplemental Figure 4).

**Figure 4.**
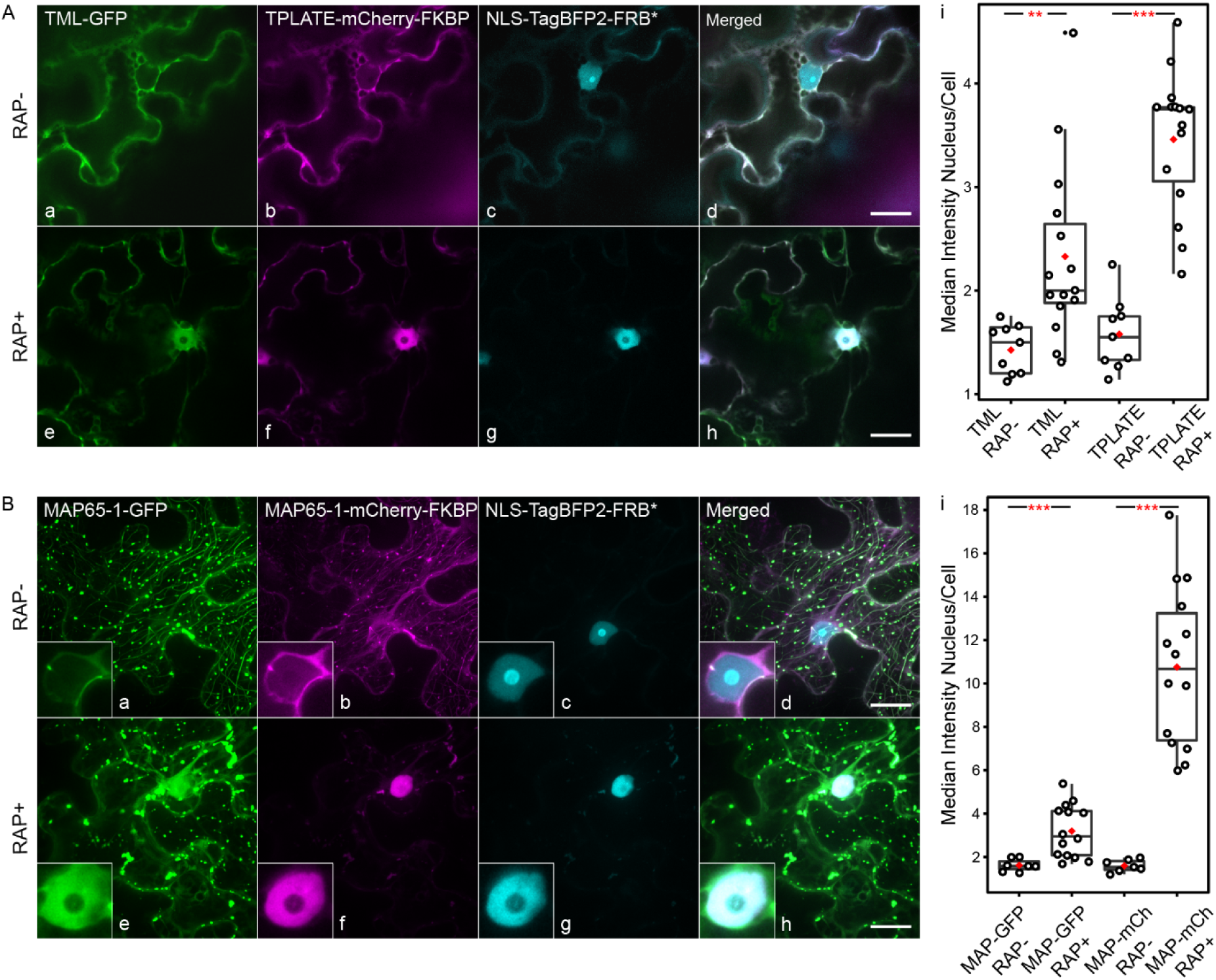
KSP allows quantitative visualization of protein-protein interactions to be performed in plants via delocalization to the nucleus. (A-B) Representative Z-stack projected images of epidermal *N. benthamiana* cells transiently expressing various GFP-fused proteins (a, e), mCherry-FKBP-fused proteins (b, f) and NLS-TagBFP2-FRB* targeted to the nuclei (c, g). Panels on the right represent merged images (d, h). In the absence of rapamycin (RAP-), localization of the GFP or mCherry-FKBP fused proteins remains unchanged irrespective of the nuclear-localized TagBFP2-FRB* construct. Rapamycin effectively relocalizes FKBP fusions to the nuclei and consequently also the interacting -GFP fused proteins. The insets represent an enlarged, nuclear area (B). Scale bars = 20μm. Statistical analysis (panels i) shows the quantification of nucleus/cytoplasm intensity ratios of co-infiltrated prey-GFP and bait-mCherry-FKBP protein fusions before and after rapamycin treatment. The black line represents the median and the red diamond represents the mean of the analyzed values. Each dot represents an individual cell. P-values <0.001 are represented as *** and <0.01 as **. Error bars represent the 95%-confidence interval.

The above examples demonstrate that KSP is inducible and feasible at various subcellular locations. This allows PPIs of different biological natures to be visualized using different subcellular anchor points. Finally, PPI can be quantified with our MitTally and NucTally scripts, which generate delocalization ratios that allow statistical analysis of the interactions to be performed.

### KSP works quickly and remains stable for several hours

To investigate the required timing for KSP to occur in *N. benthamiana*, we treated leaves co-expressing BES1-GFP, BIN2-mCherry-FKBP, and MITO-TagBFP2-FRB* with rapamycin and monitored the delocalization at different time points. Interaction through delocalization was visible as soon as five minutes after rapamycin treatment (Supplemental Figure 5A). This time is the minimal time required for preparing the sample, mounting it on the microscope, and finding the proper focal plane. To address how delocalization efficiency would proceed over time, we co-infiltrated leaves with two other protein pairs, SACL3-GFP with LHW-mCherry-FKBP and TML-GFP with TPLATE-mCherry-FKBP, together with mitochondrial or nuclear FRB* anchors, respectively. We added rapamycin and compared delocalization after 1 h and after 24 hrs in the greenhouse. SACL3/LHW rerouting to the mitochondria and TML/TPLATE rerouting to nucleus were clearly visible within one hour following treatment and remained stable over time (Supplemental Figure 5B,C). We observed a clear trend of increased delocalization over time, albeit without statistically significant differences in intensity ratios between short and long treatment for these protein pairs. These three examples indicate that KSP allows PPI to be visualized over a broad time window. Interactions occur rapidly, and at the same time they are long lasting; hence, the timing of visualization can be flexible.

### Knocksideways does not stabilize protein-protein interactions

KSP allowed us to reproduce interactions reported before (Anwar et al., 2018; De Rybel et al., 2013; Gadeyne et al., 2014; Ohashi-Ito and Bergmann, 2007; Planas-Riverola et al., 2019; Tulin et al., 2012; Vera-Sirera et al., 2015). However, we also encountered protein pairs that did not interact in our system. One canonical interaction that we tested and that does not work in our system is KIP-RELATED PROTEIN 2 (KRP2)/CELL DIVISION CONTROL 2 (CDKA;1) (Boruc et al., 2010a). We co-infiltrated *N. benthamiana* leaves with KRP2-GFP and CDKA;1-mCherry-FKBP and MITO-TagBFP2-FRB*. CDKA;1 was effectively rerouted to the mitochondria after rapamycin treatment (Figure 1D and Supplemental Figure 6A), yet KRP2-GFP remained exclusively nuclear (Supplemental Figure 6A). To test whether delocalizing KRP2 to the mitochondria (using KRP2-mCherry-FKBP) would overcome its strong nuclear targeting and therefore would be more efficient in recruiting CDKA;1-GFP, we swapped the tags on both proteins. We observed that mitochondrial targeting of KRP2 remained highly inefficient upon rapamycin treatment, even after prolonged exposure (Supplemental Figure 6B). Moreover, in those instances where we could observe KRP2 delocalization, this only rarely correlated with CDKA;1-GFP accumulation at the mitochondria (Supplemental Figure 6B f, compare the yellow arrowhead with the red arrowheads).

The PP2A subunit RCN1-mCherry-FKBP can delocalize to both the plasma membrane via MYRI-TagBFP2-FRB* (Figure 1C) and to the mitochondria with MITO-TagBFP2-FRB* (Supplemental Figure 6C). The interaction between the PP2A regulatory A subunit RCN1 and the PP2A regulatory B subunit FASS was previously shown via BiFC and tandem affinity purification (Spinner et al., 2013). Here, we were not able to replicate this interaction, as GFP-FASS was not rerouted to the FRB* anchor together with RCN1 (Supplemental Figure 6C).

The interplay between CLATHRIN LIGHT CHAIN 1 (CLC1) and AUXILIN-LIKE1 (AUXL1) was recently established *in planta* via BiFC (Adamowski et al., 2018). We co-infiltrated leaves with GFP-AUXL1 and CLC1-mCherry-FKBP and MITO-TagBFP2-FRB*. However, even though rapamycin delocalized CLC1 to the mitochondria, the localization of AUXL1 remained unaffected in our system (Supplemental Figure 6D).

At this point, it remains unclear why we could not confirm these previously reported interactions. Steric hindrance of our mCherry-FKBP tag, the requirement for a specific subcellular localization, or the requirement for additional partners for interactions to occur are likely possibilities. As our system does not stabilize PPI, KSP might also be discriminating the visualization of strong from weak or transient interactions.

### Knocksideways allows complex multi-protein interactions to be visualized *in planta*

TPLATE, TML, LOLITA, and TASH3 are part of the TPC; using one of these as bait retrieves the others as prey in Affinity Purification Mass Spectrometry (APMS) experiments (Gadeyne et al., 2014; Hirst et al., 2014). Whereas interaction between TPLATE and TML could be established using KSP (Figure 3D and Figure 4A), the smallest TPC core subunit, LOLITA-GFP, was not rerouted to the mitochondria when combined with TPLATE-mCherry-FKBP and MITO-TagBFP2-FRB* (Figure 5A). LOLITA and TASH3 were previously reported as the only interacting TPC pair in yeast (Gadeyne et al., 2014). We therefore decided to repeat the LOLITA/TPLATE interaction assay by KSP in the presence of TASH3. To avoid possible fluorophore crosstalk during spinning disk confocal microscopy, we cloned the fluorophore-less mitochondrial anchor MITO-FRB* (∼22,5 kDa). Consequently, we used KSP in a quaternary combination by co-infiltrating leaves with LOLITA-GFP, TPLATE-mCherry-FKBP, TASH3-TagBFP2, and MITO-FRB*. Without rapamycin, the fluorescently labeled proteins were omnipresent, with LOLITA also displaying a strong nuclear localization. In contrast to our previous experiment, the addition of rapamycin now rerouted all three TPC subunits to the mitochondria (Figure 5B). This indicates that all three proteins interact and that the TPLATE/LOLITA interaction requires stabilization via TASH3. We conclude that KSP allows higher-order, linear interactions to be visualized and can provide structural insights into multi-protein complexes.

**Figure 5.**
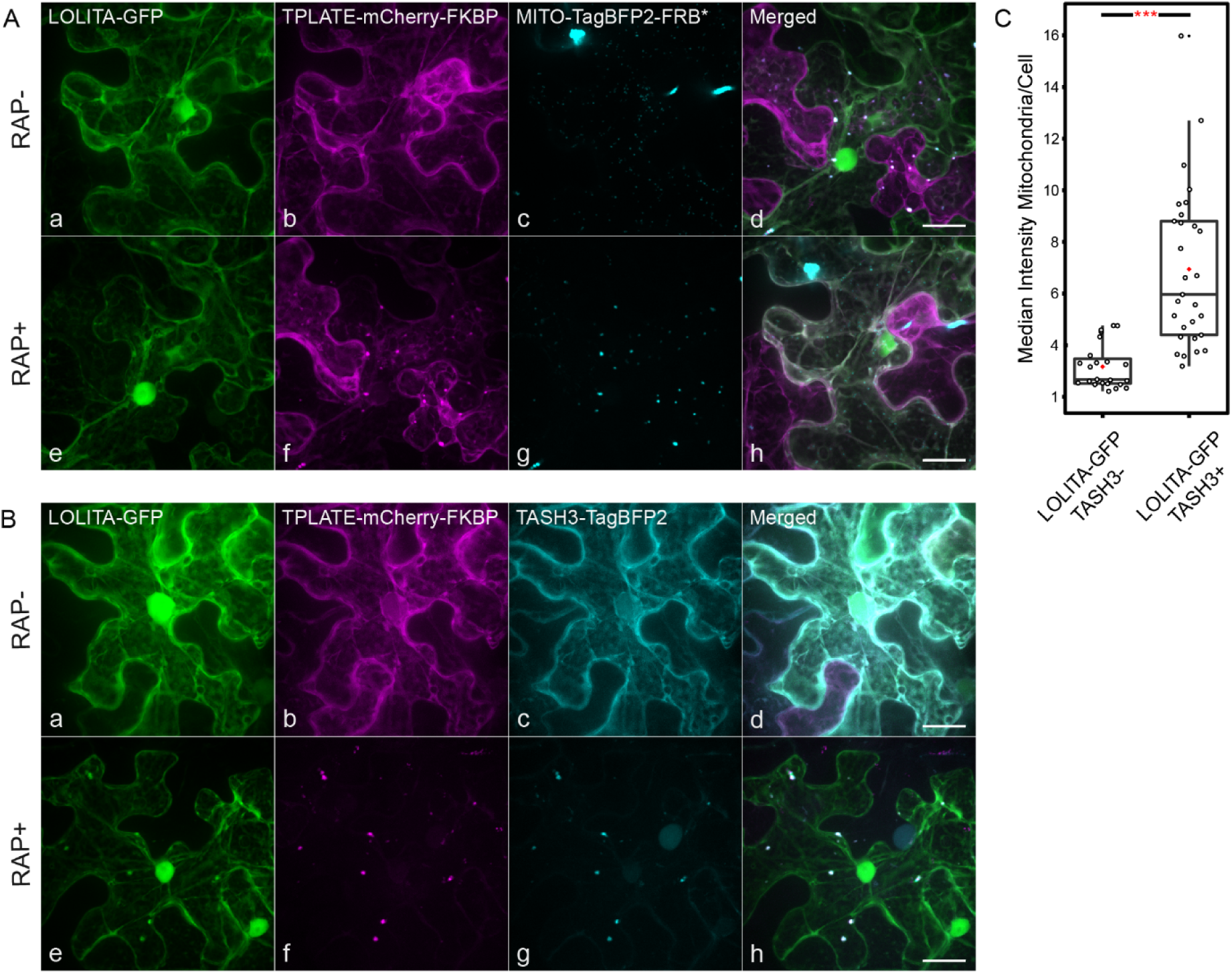
KSP allows protein-protein interactions requiring a third partner to be visualized *in planta*. (A) Representative Z-stack projections of *N. benthamiana* cells transiently expressing LOLITA-GFP (a, e), TPLATE-mCherry-FKBP (b, f), MITO-TagBFP2-FRB* (c, g) and the merged images (d, h) in the absence (RAP-) or presence (RAP+) of rapamycin. Following rapamycin treatment, TPLATE-mCherry-FKBP (f) relocalizes from the cytoplasm to the mitochondria (g), while the nuclear and cytoplasmic localization of LOLITA-GFP remains unchanged (e). (B) Representative Z-stack projections of cells transiently expressing LOLITA-GFP (a, e) TPLATE-mCherry-FKBP (b, f) TASH3-TagBFP2 (c, g) as well as MITO-FRB* without fluorescent label, in the absence (RAP-) and presence (RAP+) of rapamycin. Introduction of TASH3-TagBFP2 as a third partner stabilizes the interaction between LOLITA-GFP and TPLATE-mCherry-FKBP and allows all three proteins to relocalize to the mitochondria after rapamycin treatment (RAP+) (d, h). Scale bars = 20μm. (C) Statistical analysis showing the quantification of mitochondria/cytoplasm intensity ratios of LOLITA-GFP in the absence (left) or presence (right) of TASH3. The black line represents the median and the red diamond represents the mean of the analyzed values. Each dot represents an individual cell. P-value <0.001 is represented as ***. Error bars represent the 95%-confidence interval.

## DISCUSSION

### Rapamycin-induced FKBP-FRB heterodimerization is feasible *in planta*

Rapamycin-dependent rerouting of FKBP-tagged proteins to FRB-tagged mitochondria (KS, Knocksideways) was first used to demonstrate the role of AP-1 in retrograde trafficking in HeLa cells (Robinson et al., 2010). The FKBP-FRB-rapamycin system has so far not been widely employed in plants. This system has been used to investigate Arabidopsis cryptochrome 2 activity upon chemical induction (Rosenfeldt et al., 2008). Additionally, Arabidopsis FRB (AtFRB) and human FKBP (HsFKBP) were used to test the feasibility of the split-luciferase assay (SFLC) to monitor the dynamics of PPI in plant cells (Li et al., 2011). Compared to other fields, *in planta* tools available to investigate protein-protein interactions are relatively limited. Here, we describe KS for plants (KSP) in the *N. benthamiana* transient expression assay, which we developed into an inducible PPI tool holding a unique set of features.

We show that rerouting various proteins to distinct subcellular localizations is feasible in our system. Other ‘anchor-away’ techniques such as NTA, CAPPI and MILo (Dixon and Lim, 2010; Kaplan--Levy et al., 2014; Lv et al., 2017) are limited in terms of subcellular rerouting options due to their direct fusion with the anchor. By contrast, KSP easily combines with multiple anchors without the need to re-clone fusion constructs. In the current study, we designed four different anchors, yet hypothetically our system is compatible with any other desired subcellular targeting (Nelson et al., 2007). We tested various FKBP-tagged bait proteins of interest (POIs) and successfully delocalized all of them to microtubules, to the plasma membrane, to the mitochondria, and to the nucleus. We showed that rapamycin-dependent FKBP-FRB* heterodimerization is robust and capable of trapping POIs at an ectopic location. Also, rerouting is likely to occur more rapidly than targeting to a specific location by conditionally expressing nanobodies (Früholz et al., 2018), as our approach does not rely on any translational step to achieve the delocalization. We observed rerouting of POIs in *N. benthamiana* within minutes, which is in agreement with very fast kinetics of FKBP-FRB binding upon rapamycin induction in *Arabidopsis* reported previously (Li et al., 2011).

### KSP is a valuable tool for visualizing PPIs

We thoroughly investigated previously reported PPIs by KSP using mitochondrial and nuclear anchors. We demonstrated that our tool could successfully be used to identify interactions between proteins with diverse physiological roles in distinct biological processes. We successfully reproduced previously demonstrated interactions of the transcription factors SACL3 and LHW (Vera-Sirera et al., 2015), the interplay of TPLATE and TML, subunits of endocytic TPLATE complex (Gadeyne et al., 2014), and interaction between the Brassinosteroid signaling proteins BIN2 and BES1 (Anwar et al., 2018; Planas-Riverola et al., 2019). Likewise, the homodimerization of MAP65-1 (Smertenko et al., 2004; Tulin et al., 2012; Van Damme et al., 2004) was successfully identified using our system.

At the same time, using KSP, PPIs are visualized only upon drug exposure, which we consider one of the major advantages of our system, as this allows additional control infiltrations to be omitted. Rapamycin-induced visualization of interactions is at the same time rapid and long lasting. This allows POIs to be monitored throughout the entire experiment, and the visualization of PPIs can be induced by drug treatment at any desired time point. The high affinity of rapamycin dimerization (Banaszynski et al., 2005), however, makes it difficult to reverse. Ascomycin, a rapamycin competitor for HsFKBP (Paulmurugan et al., 2004), was shown to work in plant protoplasts (Li et al., 2011). We therefore investigated the capacity of the rapamycin competitor ascomycin to reverse the delocalization. We showed that the FKBP-FRB* dimerization is partially reversible in our system, albeit it requires prolonged treatment. We speculate that in *N. benthamiana*, syringe-based administration of the drug is difficult to control when substituting rapamycin with its competitor. It also remains possible that the efficiency of ascomycin depends on the pair of the POIs tested, as well as their subcellular location.

Another advantage of KSP is its straightforward readout with reduced dependence on fluorophore choice and/or proximity of the bait and prey fluorophores. This is quite different from BiFC or FRET-FLIM assays, where spatial tagging has a very prominent effect on the outcome of the interaction readout. In addition, in contrast to BiFC, KSP is less prone to false positive results, which can be generated by (for example) unstable proteins. False negatives, however, caused by instability of the FKBP-fused bait protein remain possible. To rule out this possibility, the expression of full-length fusions by immunoblotting can be tested.

Biological processes are regulated by many mechanisms, including higher-order cooperation between proteins. Evaluating the interplay among multimeric proteins remains a challenging task. Three-fluorophore FRET-FLIM and trimolecular fluorescence complementation (Glöckner et al., 2019; Offenborn et al., 2015) have been shown to work in plants. However, the requirement for spatial tagging in order to obtain a positive readout using the above tools likely limits the analysis of any random protein complex of choice. By contrast, our KSP system holds great potential for visualizing higher-order PPIs, which we were able to demonstrate for three members of the endocytic TPC. Our system is theoretically compatible with even higher numbers of interactions and could provide insight into the structural properties of TPC or other complexes.

Another great advantage of KSP lies in its quantification capabilities. We developed semi-automatic quantification scripts for mitochondrial (MitTally) and nuclear (NucTally) anchors. These scripts produce straightforward results, which are expressed as the intensity ratio between particles (mitochondria or nuclei) and the cytoplasm. In addition, they provide both raw data results and results with outliers excluded via the interquartile range method. Both scripts require only moderate effort, as images can be analyzed in batches, and the results are easy to subject to statistical processing.

The limitations of the KSP system we encountered are as follows: weak and transient interactions might not be efficiently detected; and the tool does not work if the prey itself is anchored to a specific location or if the prey is part of a stronger interaction that anchors it, as our bait and prey proteins are not artificially coupled. The observed lack of interaction between CDKA;1 and KRP2, regardless of the orientation of the - GFP and -mCherry-FKBP tagging, might be due to the strong nuclear anchoring of KRP2 and as well as the requirement for another interacting partner, likely a cyclinD (Zhou et al., 2002). Establishing spatial targeting inside the nucleus might help to solve this issue. TPX-Like proteins were recently shown to cause intranuclear MT polymerization *in planta* (Boruc et al., 2019); therefore, TPX-Likes could potentially be used in KSP as FRB* intranuclear anchors to visualize specific nuclear PPIs (Boruc et al., 2010b).

### Future perspectives

We demonstrated that KSP can be used to visualize multiple proteins interactions. A logical next step would be to increase the number of interactors tested in a single assay, especially since the imaging of multiple fluorophores expressed together is becoming easier in the context of novel technical advances in confocal microscopy (Borlinghaus et al., 2006; Min et al., 2014). However, infiltrating multiple constructs into *N. benthamiana* leaves may lead to uneven expression patterns and ratios of POIs. One solution would be to combine the expression of multiple proteins from a single vector to ensure the same expression levels of all of the co-expressed elements (Decaestecker et al., 2019; Engler et al., 2008; Sarrion-Perdigones et al., 2011).

KSP also has the potential to serve as an interactomics tool. Trapping FKBP-fused bait proteins and their interactors at the mitochondria could be combined with mitochondrial fraction isolation. Comparative mass spectrometry analysis of rapamycin-treated versus untreated samples will allow specific interacting partners to be analyzed. KS was recently combined with BioID into the two-component (2C)-BioID system to conditionally target the biotin ligase to a specific subcellular location in human fibroblast cells (Chojnowski et al., 2018). As BioID has recently been adapted for plants (Arora et al., 2020; Mair et al., 2019; Zhang et al., 2019), 2C-BioID could also be introduced *in planta* in order to improve the overall robustness of the assay.

As we have also shown here by targeting a phosphatase affecting PI4P levels to the PM, KSP can also be further developed beyond a tool for detecting PPIs. Introducing KSP into other model systems such as *Arabidopsis thaliana* will create the possibility for conditional targeting of proteins and their interaction partners to a subcellular location. In animal cells, rapamycin-induced PM targeting of phosphatidylinositol phosphatases or rerouting AP-2 were used to acutely control endocytosis (Hammond et al., 2012; Heo et al., 2006; Wood et al., 2017). Possible applications in plants might therefore also include the acute and conditional targeting of proteases, kinases, or phosphatases to a specific subcellular location.

Similar to KS in animals (Robinson et al., 2010), KSP might become a conditional *in planta* knockout tool, which could be used to study otherwise lethal mutants (Lloyd and Meinke, 2012; Lloyd et al., 2015). Moreover, combining KSP with tissue-specific promoters (Siligato et al., 2016) would allow live-imaging to study the immediate effects of inducible protein knockouts in particular plant cell types, tissues, and organs while avoiding the pleiotropic effects of gene loss using conventional approaches. This approach might overcome limitations in terms of capacity and/or timing observed by conditionally delocalizing functional proteins using nanobody expression (Winkler et al., 2020). Rapamycin is not a very potent inhibitor of the plant Target of Rapamycin (TOR) kinase, which controls growth in response to environmental signals. However, over-expression of a non-plant FKBP12 enhances the inhibitory activity of rapamycin towards TOR (Ren et al., 2012; Sormani et al., 2007). The use of KSP in Arabidopsis to investigate developmental or physiological effects will therefore always have to take into account possible side-effects of inhibiting the TOR kinase pathway to some extent. Optimal KSP experiments in Arabidopsis should therefore use FKBP constructs expressed at low levels in combination with strict controls (i.e. the use of plants expressing similar levels of the FKBP-fusions in combination with rapamycin treatment, but without the FRB* anchor).

Taken together, our results demonstrate that KSP is robust and efficient tool for PPI visualization in plants. As KSP holds specific advantages and can visualize complex multi-protein interactions, it will likely open a new path to investigate protein-protein interactions in plants complementary to the currently existing tools.

## METHODS

### Multisite Gateway cloning of entry clones

Gateway entry clones pDONR221-TagBFP2, pDONR221-MITOTagBFP2, pDONR221MITO, pDONR221-MYRI, pDONR221-LHW, pDONR221-SACL3, pDONR221-FKBP-mCherry, pDONRP2RP3-mCherry-FKBP, pDONRP2RP3-FRB* and pENTR™ 5’-TOPO entry clone pUBQ10NLS were generated in this study according to the manufacturer’s instructions (ThermoFisher Scientific BP Clonase). pDONR221-TagBFP2 was amplified from pSN.5 mTagBFP2 (Pasin et al., 2014). pDONR221-MITOTagBFP2 was generated from pDONR221-TagBFP2 by including the import signal of the yeast mitochondrial outer membrane protein Tom70p as described before (Robinson et al., 2010). pDONR221-MITO was generated from pDONR221-MITOTagBFP2. pDONR221-MYRI, consisting of the N-terminal fragment (amino acids 1 to 76) of CPK34, which contains a myristoylation/palmitoylation site (Kirik et al., 2012; Myers et al., 2009; Podell and Gribskov, 2004), was synthesized by Gen9 (now Ginkgo Bioworks, 27 Drydock Avenue, Boston, MA 02210). pDONR221-LHW and pDONR221-SACL3 were kindly provided by Prof. Bert De Rybel (PSB, Ghent, Belgium). pDONR221-FKBP-mCherry was generated by sewing PCR with specific primers. The first fragment contained the AttB1 and FKBP-linker sequences and the second fragment contained the linker-mCherry and AttB2 sequences, which were then fused via overhang PCR followed by BP cloning. pDONRP2RP3-mCherry-FKBP was generated by sewing PCR. The first PCR was performed with mCherry-specific primers (Mylle et al., 2013). The forward primer contained the AttB2r site and the N-terminal mCherry sequence, and the reverse primer contained the C-terminal mCherry sequence and the linker. The second PCR was performed with FKBP-specific domain (Robinson et al., 2010) primers. The forward primer contained the linker sequence and the N-terminal FKBP sequence and the reverse primer contained the C-terminal FKBP sequence flanked with AttB3. pDONRP2RP3-FRB* was amplified from MAPPER (Chang et al., 2013, 2017) with FRB* domain-specific primers flanked with AttB2r and AttB3 sites. Star “*” indicates the threonine 2098 to leucine mutation, which allows rapamycin to be replaced with its analogs, rapalogs (Bayle et al., 2006). pUBQ10NLS was generated using pUBQ forward and NLS reverse primers using the pBINU-NYA(K) (N799872) plasmid as template (Mehlmer et al., 2012). The overhangs were generated using Taq polymerase, and the purified amplicon was ligated into pENTR 5’TOPO (ThermoFisher Scientific) according to the manufacturer’s instructions. The following entry clones were described before: pDONR207-EB1A, pDONR207-MAP65-1, pDONR207-T22.1/TPLATE (Van Damme et al., 2004), pDONR207-CLC1 At2g20760 (Van Damme et al., 2011), pDONR207-RCN1/PP2AA1 (Spinner et al., 2013), pDONR221-CDKA;1 (Boruc et al., 2010b), pDONR221-TASH3 (Gadeyne et al., 2014), pDONR221-BIN2, (Houbaert et al., 2018), pDONRP2RP3-Sac1 and pDONRP2RP3Sac1-dead (Doumane and Caillaud, 2020). All entry clones used in this study are listed in Supplemental Table 1. All primers sequences are shown in Supplemental Table 1. Schematic representations of entry clones created *de novo* in this study are shown in Supplemental Figure 1.

### Cloning of LR binary vectors

Gateway expression clones were obtained after LR recombination using LR Clonase according to the manufacturer’s instructions (Thermo Fisher). CDKA;1-mCherry-FKBP, CLC2-mCherry-FKBP, EB1A-TagBFP-FRB, FKBP-mCherry-Sac1, FKBP-mCherry-Sac1-dead, KRP2-mCherry-FKBP, LHW-mCherry-FKBP, MAP-65-1-mCherry-FKBP, MITOTagBFP2-FRB, MYRI-TagBFP2FRB, pUBQ10NLS-TagBFP2-FRB, RCN1/PP2AA1-mCherry-FKBP, SACL3-GFP, TASH3-TagBFP2, and TPLATE-mCherry-FKBP were cloned in pB7m34GW and under the control of the p35S promoter, except for MITOTagBFP2-FRB* and pUBQ10NLS-TagBFP2-FRB*, which were cloned under the control of the pH3.3 (Ingouff et al., 2017) and pUBQ (Mehlmer et al., 2012) promoters, respectively. The eGFP sequence was cloned into pK7WG2 (Karimi et al., 2002, 2005). The following expression clones were described before: pB7m34GW-pUBQ:GFP-AUXILIN-LIKE1 (Adamowski et al., 2018), pGWB6-GFP-FASS/TON2 (Spinner et al., 2013), pK7WG2-p35S:KRP2-GFP, pK7WG2-p35S:CDKA;1-GFP (Boruc et al., 2010b), pK7FWG2-35S:TML-GFP, pK7FWG2-p35S:LOLITA-GFP (Gadeyne et al., 2014), pK7FWG2-p35S:MAP65-1-GFP (Van Damme et al., 2004), UBQ10prom:mCitrine-P4M (Doumane and Caillaud, 2020; Simon et al., 2016). 35S:BES1-GFP was a kind gift from Prof. Jenny Russinova (PSB, Ghent, Belgium). All expression vectors and accession numbers are presented in Supplemental Table 2.

### Plant growth and transient expression assay

*Nicotiana benthamiana* plants were grown in a greenhouse under long-day conditions (06 h to 22 h light, 100 PAR, 21 °C) in soil (Saniflo Osmocote pro NPK: 16-11-10 + Magnesium and trace elements). Transient expression was performed by leaf infiltration (Sparkes et al., 2006) with some modifications. *Agrobacterium* OD of MITOTagBFP2-FRB was adjusted to 0.1 prior to mixing the cultures to assure sufficient protein expression while avoiding massive mitochondrial clustering. *N. benthamiana* leaves were imaged two days after infiltration. Imaging was performed on a PerkinElmer Ultraview spinning-disc system attached to a Nikon Ti inverted microscope and operated using the Volocity software package. Images were acquired on an ImagEM CCD camera (Hamamatsu C9100-13) using frame-sequential imaging with a 60x water immersion objective (NA = 1.20). Specific excitation and emission was performed using a 405nm laser excitation combined with a single band pass filter (454-496nm) for TagBFP2, 488nm laser combined with a single band pass filter (500-550nm) for GFP, 514nm laser excitation combined with single band pass filter (550nm) for mCitrine and 561nm laser excitation combined with a dual band pass filter (500-530nm and 570-625nm) for mCherry. Images shown are Z-stack projections, except the insets, which represent enlarged, single slices (Figure 1A and B, Figure 3A, Figure 4B, Supplemental Figure 4). Z-stacks were acquired in sequential frame mode at a 1 µm interval using the Ultraview (Piezo) focus drive module. Cells in which the -FKBP protein fusions were not rerouted to the FRB* anchors after rapamycin treatment were not imaged.

### Drug and mitochondrial marker treatments

*N. benthamiana* leaves at 48 h after infiltration were treated with 1 µM rapamycin (Sigma-Aldrich) and/or 50 µM oryzalin (Sigma-Aldrich) or 250 nM MitoTracker™ Red CMXRos (Thermo Fisher) and/or 100 µM ascomycin (MedChemExpress) by syringe-infiltration. Stock solutions (1 mM rapamycin, 20 mM oryzalin, 1 mM MitoTracker and 10 mM ascomycin) were prepared by diluting the corresponding amount of chemical in DMSO (dimethyl sulfoxide). Prior to infiltration, the chemicals were diluted to their final concentrations in MiliQ water.

### Quantification of delocalization ratio and statistical analysis

Raw pictures (16bit gray scale) in Figure 1, 2, 3, 4, 5, Supplemental Figure 2, Supplemental Figure 3, Supplemental Figure 4, Supplemental Figure 5 and Supplemental Figure 6 were exported from Volocity (Perkin Elmer) as single files with merged Z-planes and separate channels. Pictures were processed in ImageJ/Fiji (Schindelin et al., 2012) as described below (see Supplemental File 1). Mitochondria or nuclear vs. cytoplasm intensity ratios were generated with MitTally (Figure 3 via version 10, Figure 5 and Supplemental Figure 5 via version 13) or NucTally (Figure 4 via version 13 and Supplemental Figure 3 and Supplemental Figure 5 via version 17) and were analyzed in RStudio (Version 1.2.5033) with R (RStudio, 2015). For mitochondrial rerouting analysis, outliers were automatically removed by MitTally based on particle median intensities in a single step interquartile computation. For nuclear rerouting analysis, outliers were removed by hand based on average N/C median ratios in a single-step interquartile computation (Figure 4) or automatically by NucTally based on N/C ratios calculated from median intensity values via single-step interquartile computation (Supplemental Figure 3 and Supplemental Figure 5). The number of endosomes in Figure 2 was counted manually and values were normalized to the selected cell area. Statistical analysis in Figures 2-5 and Supplemental Figure 3 was done with ANOVA to account for heteroscedasticity. Post hoc pairwise comparison was performed with the package MULTCOMP utilizing the Tukey contrasts (Herberich et al., 2010). Group characterization in Supplemental Figure 3F was done by transforming p-values with the multicompView package. Statistical analysis in Supplemental Figure 5 was performed using the Mann-Whitney-Wilcoxon test. Tables with the results of the statistical analysis can be found in Supplemental File 4. The most recent versions of the scripts, MitTally (version 13; Supplemental File 2 and NucTally (version 17; Supplemental File 3), as well as a detailed manual to use these scripts (Supplemental File 1), are available as supplemental data.

### Accession Numbers

The Arabidopsis Information Resource (TAIR) locus identifiers for the genes mentioned in this study are AUXILIN-LIKE1 (At4G12780), BES1 (At1G19350), BIN2 (At4G18710), CDKA;1 (At3G48750), CLC1 (At2G20760), EB1a (At3G47690), FASS/ TON2 (At5G18580), KRP2 (At3G50630), LHW (At2G27230), LOLITA (At1G15370), MAP65-1 (At5G55230), MYRI/CPK34 (At5G19360), RCN1/PP2AA1 (At1G25490), SACL3 (At1G29950), T22.1/TPLATE (At3G01780), TASH3 (At2G07360), TML (At5G57460). MitTally and NucTally scripts were uploaded to the GitHub platform and are available via the following link: https://github.com/winkler-joanna/MitTally_NucTally.git

## Supplemental Data

**Supplemental Figure 1. Schematic representation of the available Multisite Gateway entry clones and expression constructs used in this study**.

**Supplemental Figure 2**. MITO-TagBFP2-FRB* occasionally causes mitochondrial clustering without interfering with their capacity to recruit FKBP-fused proteins.

**Supplemental Figure 3**. FKBP-FRB* heterodimerization is partially reversible via prolonged ascomycin treatment.

**Supplemental Figure 4**. Microtubule depolymerization does not relocalize MAP65-1 to the nuclei.

**Supplemental Figure 5**. Rapamycin treatment relocalizes interacting proteins rapidly and remains effective over long periods.

**Supplemental Figure 6**. KSP does not stabilize protein-protein interactions and visualizing interactions via delocalization therefore has some limitations.

**Supplemental Table 1**. List of primers used in this study.

**Supplemental Table 2**. List of constructs used in this study.

**Supplemental File 1**. Manual of the quantification scripts.

**Supplemental File 2**. MitTallyV13 script.

**Supplemental File 3**. NucTallyV17 script.

**Supplemental File 4**. Statistical analysis.

## ACKNOWLEDGMENTS

The authors would like to thank Prof. Jiri Friml (IST, Klosterneuburg, Austria) and Dr. Sibu Simon (CEITEC, Brno, Czech Republic) for the help in initiating this project and many colleagues who have provided material for this work. pSN.5 mTagBFP2 was kindly provided by Dr. Fabio Pasin (Academia Sinica, Taipei, Taiwan). pDONR221-LHW and pDONR221-SACL3 were kindly provided by Prof. Bert De Rybel (PSB, Ghent, Belgium). The FKBP construct was kindly provided by Prof. Margaret Robinson (CIMR, Cambridge, UK). pB7m34GW-pUBQ:GFP-AUXILIN-LIKE1 was kindly provided by Dr. Maciek Adamowski (IST, Klosterneuburg, Austria). pGWB6-GFP-FASS/TON2 was kindly provided by Dr. Martine Pastuglia (IJPPB Versailles, France). pDONR221-BIN2 and 35S:BES1-GFP were kindly provided by Prof. Jenny Russinova (PSB, Ghent, Belgium). UBQ10prom:mCitrine-P4M, pDONRP2RP3-Sac1 and pDONRP2RP3-Sac1-dead were kindly provided by Marie-Cécile Caillaud (ENS Lyon, France). Research in the Van Damme lab is supported by the European Research Council T-Rex project number 682436 and by the National Science Foundation Flanders (FWO; G009415N).

## AUTHOR CONTRIBUTIONS

DVD initiated the project and designed experiments. JW, EM, ADM and PG designed and performed experiments. BP wrote the script for quantification of the data. JM performed experiments. JW, PG and DVD wrote the paper.

The authors declare no competing interests.

## Notes

### Competing Interest Statement

The authors have declared no competing interest.

### Summary of Updates

This is an updated version of the manuscript which is now accepted at The Plant Cell.

