## supplemental data (figures and tables) for "Visualizing protein-protein interactions in plants by rapamycin-dependent delocalization"

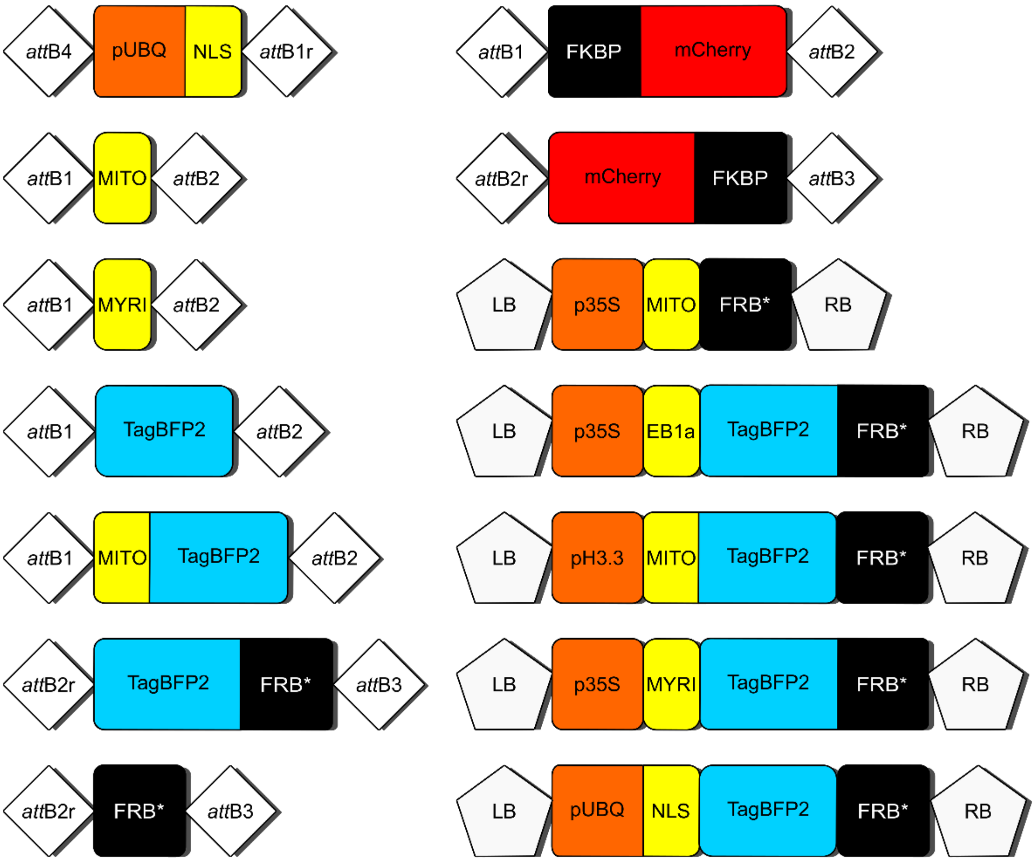


**Supplemental Figure 1. Schematic representation of the available Multisite Gateway entry clones and expression constructs used in this study.**

Abbreviations are as follows: pUBQ, ubiquitin promoter; NLS, nuclear localization signal; MITO, mitochondrial outer membrane localization signal from *Saccharomyces cerevisiae* protein Tom70p (Robinson et al., 2010); MYRI, myristoylation site, plasma membrane localization signal (Kirik et al., 2012); FRB*, FRB-domain of mammalian TOR kinase with T>L mutation (Bayle et al., 2006); FKBP, FKBP domain of HsFKBP12; EB1a, *Arabidopsis thaliana* MICROTUBULE END BINDING PROTEIN 1a (Van Damme et al., 2004); p35S, 35S promoter; pH3.3, histone H3 promoter; LB, left border of expression construct; RB, right border of expression construct. Domains are not drawn to scale. Molecular weight of KSP tags is as follows: -TagBFP2-FRB* ~84 kDa; -FRB* ~22.5 kDa; -mCherry-FKBP and -FKBP-mCherry ~86 kDa. This figure provides an overview of the available new building blocks for using KSP.

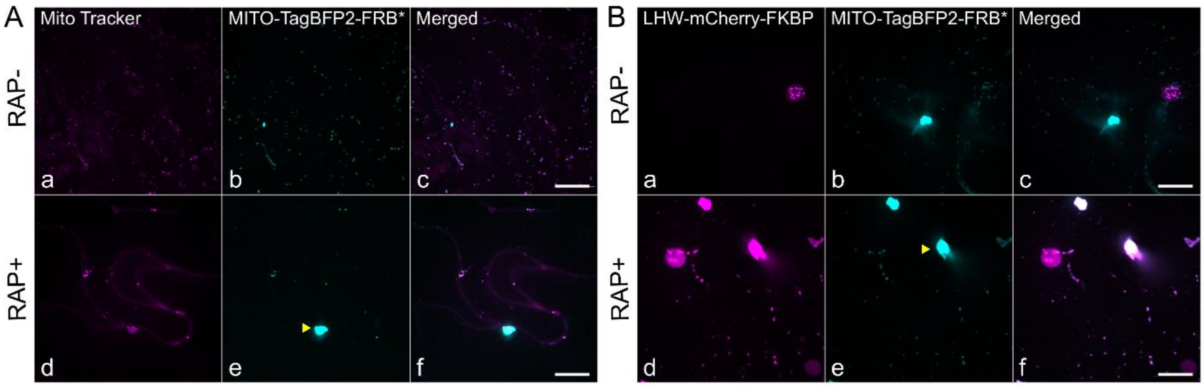


**Supplemental Figure 2: MITO-TagBFP2-FRB* occasionally causes mitochondrial clustering without interfering with their capacity to recruit FKBP-fused proteins. (Supports Figure 1, Figure 3 and Figure 5).**

(A) Representative Z-stack projections of *N. benthamiana* cells expressing MITO-TagBFP2-FRB* co-stained with the mitochondrial marker MitoTracker Red. MitoTracker Red effectively co-localizes with the MITO-TagBFP2-FRB* positive puncta, which occasionally can be seen as large clusters (yellow arrowhead). (B) Representative Z-stack projections of cells expressing LHW-mCherry-FKBP, which localizes to the nucleus (a, d) and mitochondrial-targeted, partially clustered MITO-TagBFP2-FRB* (b, e). Merged panels are shown on the right (c, f). In the absence of rapamycin (RAP-) mitochondria clustering does not alter the localization of LHW-mCherry-FKBP. After rapamycin treatment (RAP+) MITO-TagBFP2-FRB*-positive mitochondrial clusters (yellow arrowhead) effectively recruit LHW-mCherry-FKBP. Scale bars = 20µm.

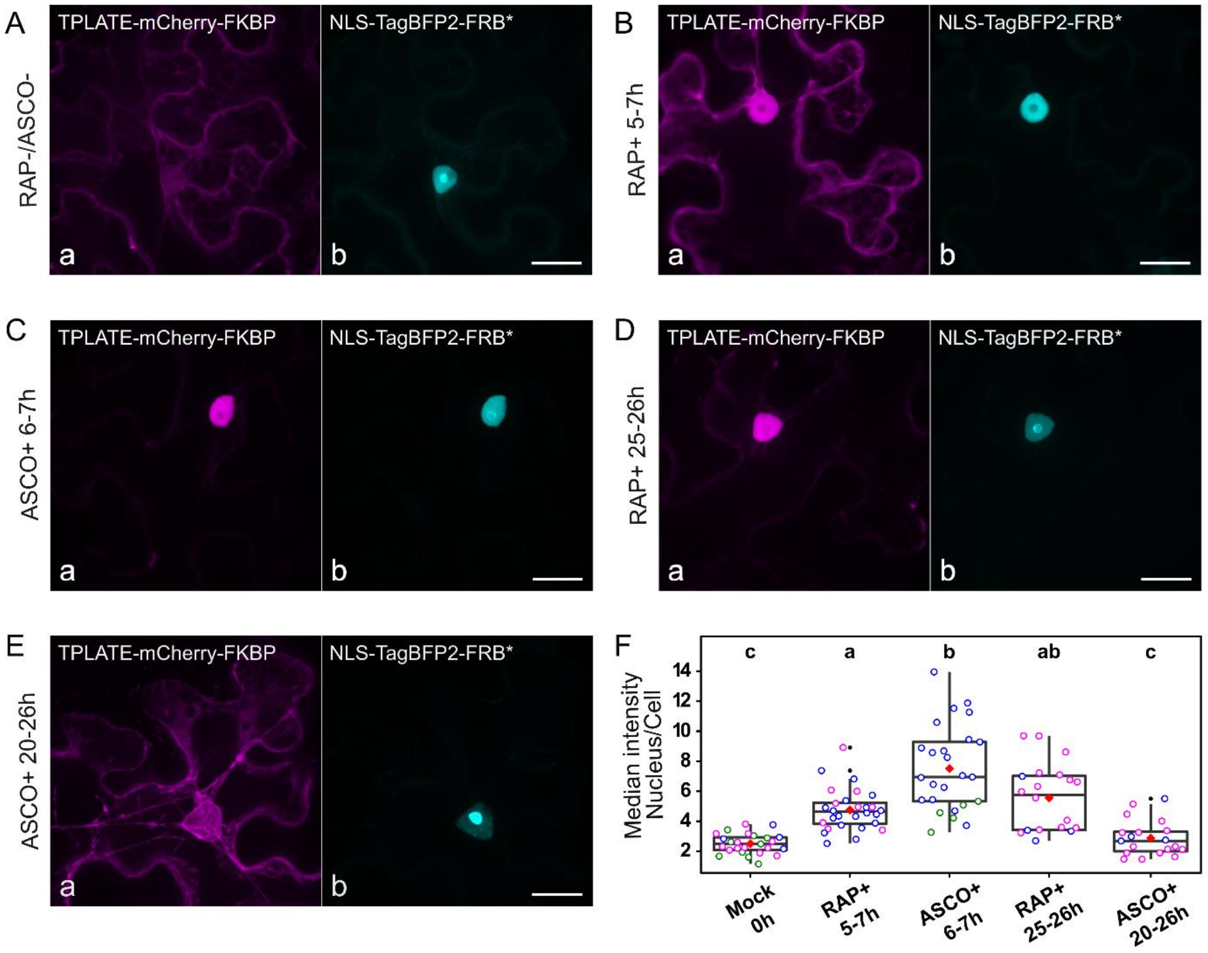


**Supplemental Figure 3: FKBP-FRB* heterodimerization is partially reversible via prolonged ascomycin treatment. (Supports Figure 1).**

(A) Without rapamycin treatment, TPLATE-mCherry-FKBP (a) mainly localizes to the cytoplasm. (B) Rapamycin treatment (RAP+) delocalizes TPLATE to the nuclear anchor NLS-TagBFP2-FRB (b) and binding remains stable over several hours. (C) In the samples, where one hour after rapamycin treatment, its competitor, ascomycin, was subsequently infiltrated, FKBP-FRB* heterodimerization was not reversed after several hours. (D) Prolonged exposure to rapamycin retains TPLATE in the nucleus. (E) In ascomycin-treated samples, prolonged treatment releases TPLATE again to the cytoplasm. Scale bars equal 20µm. (F) Statistical analysis showing the quantification of nucleus/cytoplasm intensity ratios of TPLATE-GFP under different drug exposure conditions. Significant differences between the groups are annotated by letters a-c. Notably, there is no significant difference between the Mock sample and 20-26h ASCO+ sample. The black line represents the median and the red diamond represents the mean of the analyzed values. Each dot represents an individual cell, and the different colors of the dots represent individual infiltrations. Error bars represent the 95%-confidence interval. This figure demonstrates that KSP is partially reversible.

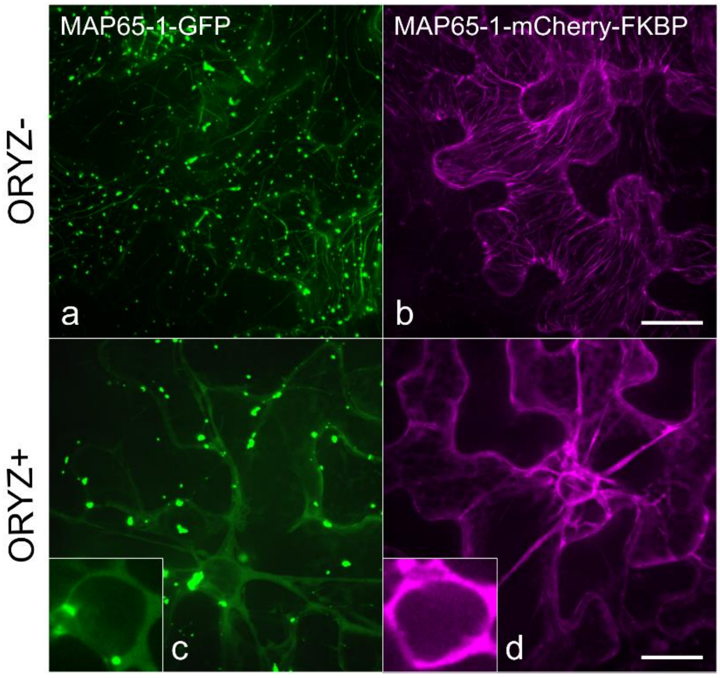


**Supplemental Figure 4: Microtubule depolymerization does not relocalize MAP65-1 to the nuclei. (Supports Figure 4).**

MAP65-1-GFP (a) and MAP65-1-mCherry-FKBP (b) localize to cortical microtubules. MAP65-1-GFP, much more than MAP65-1-mCherry-FKBP, is also visible in aggregates. The reason why the GFP-fused form of MAP65-1 clusters so heavily compared to the mCherry-fused form is unclear as both are expressed from the same promoter. Oryzalin (ORYZ+) depolymerizes microtubules (bottom row) and releases MAP65-1 to the cytoplasm. MAP65-1 remains excluded from the nuclei (c, d). Insets represent enlarged, nuclei areas. Scale bars = 20µm.

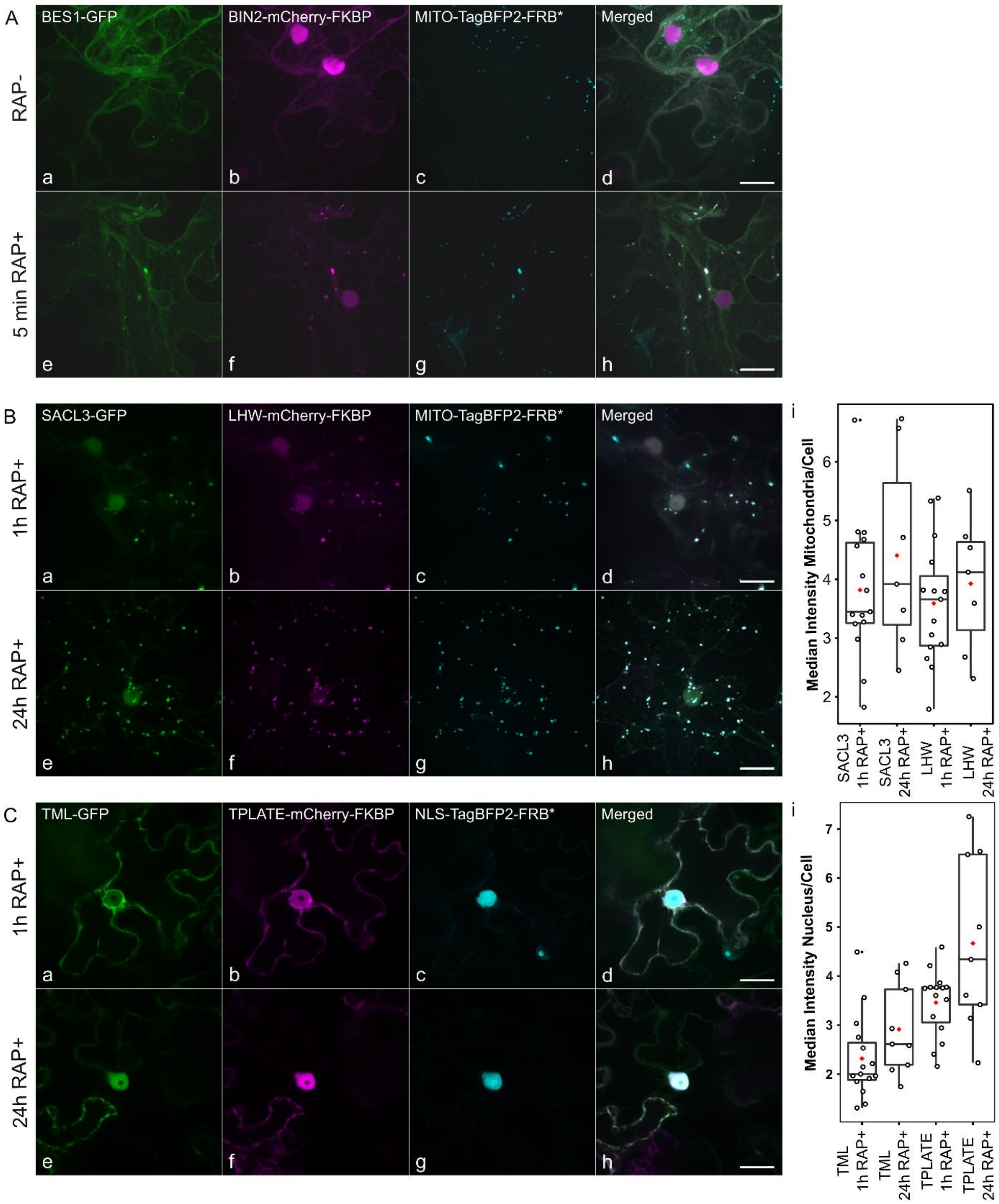


**Supplemental Figure 5: Rapamycin treatment relocalizes interacting proteins rapidly and remains effective over long periods. (Supports Figure 3 and 4).**

(A) BES1-GFP (a, e), BIN2-mCherry-FKBP (b, f), MITO-TagBFP2-FRB* (c, g), and merged images (d, h), untreated (RAP-) and treated (RAP+). Rapamycin relocalizes BES1-GFP and BIN2-mCherry-FKBP to the mitochondria within 5 minutes following the treatment. (B and C) SACL3-GFP (a and e), LHW-mCherry-FKBP (b and f), MITO-TagBFP2-FRB* (c and g) and merged images (d and h) and TML-GFP (a and e), TPLATE-mCherry-FKBP (b and f), NLS-TagBFP2-FRB* (c and g), and merged images (d and h) during short (1h RAP+) or prolonged (24h RAP+) rapamycin treatment. Rapamycin-dependent relocalization of -GFP and -mCherry-FKBP protein fusions to organelles can be visualized after 1 hour and remains stable even after 24 hours following treatment. Scale bars equal 20µm. Statistical analyses

(i) showing the quantification of intensity -GFP and -mCherry-FKBP-fused proteins, after short (1 h), and long (24 h) rapamycin treatment. Black lines represent the median and red diamonds represent the mean of the analyzed values. Each dot represents an individual cell. Error bars represent the 95%-confidence interval. Although there is a trend towards increased accumulation over time, no significant difference could be found.

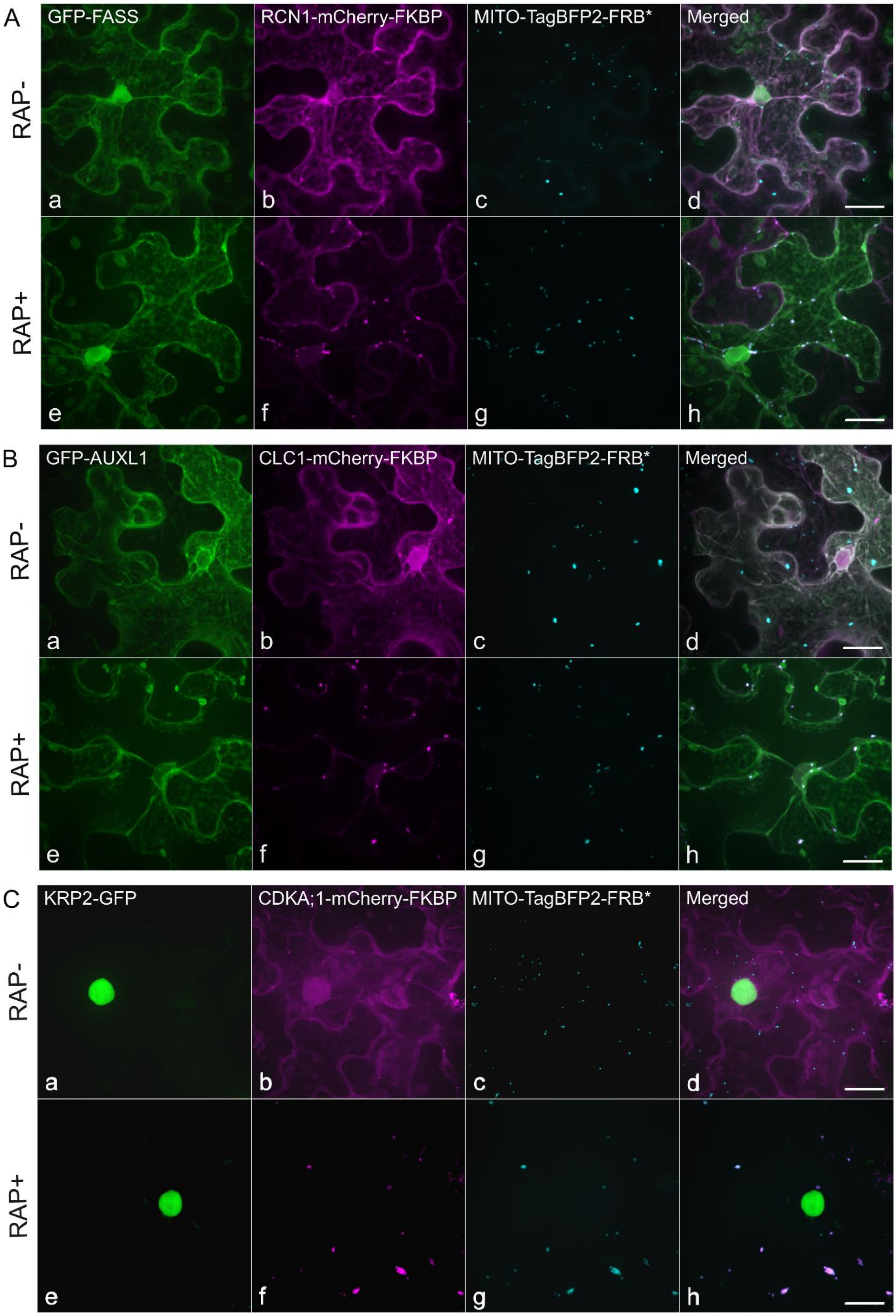
A
RAP-
RAP+
GFP-FASS
RCN1-mCherry-FKBP
MITO-TagBFP2-FRB*
Merged
a
b
c
d
e
f
g
h
B
RAP-
RAP+
GFP-AUXL1
CLC1-mCherry-FKBP
MITO-TagBFP2-FRB*
Merged
a
b
c
d
e
f
g
h
C
RAP-
RAP+
KRP2-GFP
CDKA;1-mCherry-FKBP
MITO-TagBFP2-FRB*
Merged
a
b
c
d
e
f
g
h

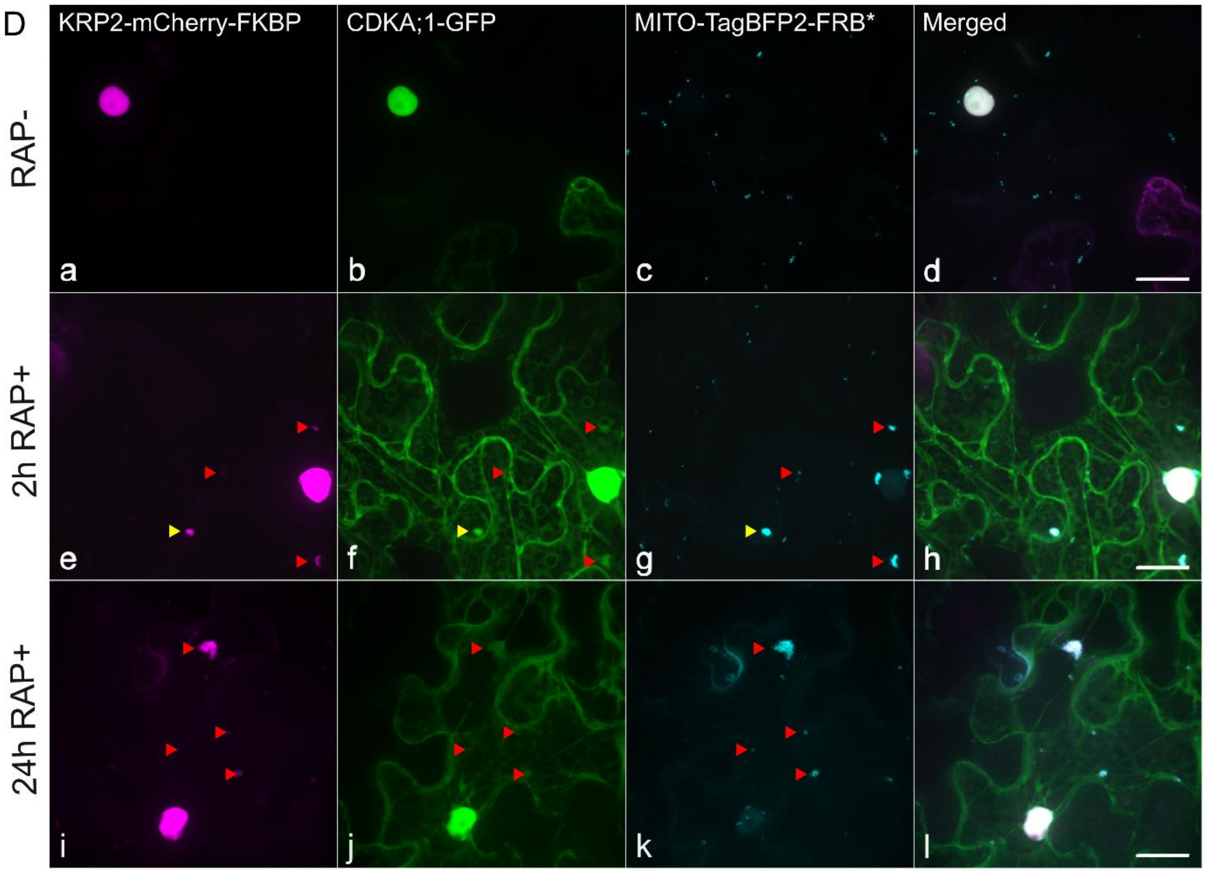


**Supplemental Figure 6: KSP does not stabilize protein-protein interactions and visualizing interactions via delocalization therefore has some limitations. (Supports Figure 3 and 4).**

(A-D) Cells transiently expressing GFP-tagged proteins (a, e), mCherry-FKBP-tagged proteins (b, f) as well as the mitochondrial anchor MITO-TagBFP2-FRB* (c, g), in the absence (RAP-) and presence (RAP+) of rapamycin. Merged panels are shown on the right (d, h). The protein pairs FASS and RCN1 (A), AUXILLIN-LIKE1 and CLC1 (B) and KRP2 and CDKA;1 (C and D) were shown previously to interact by alternative interaction assays that stabilized their interaction. After rapamycin treatment (RAP+), the FKBP-fused proteins (RCN1, CLC1 and CDKA;1) were efficiently targeted to the mitochondria. The GFP-fused (FASS, AUXL1 and KRP2) localizations remained however unchanged. To test whether the negative interaction between KRP2 and CDKA;1 could be caused by a strong nuclear retention of KRP2, we also tested the CDKA;1-KRP2 interaction in the other direction, by pulling KRP2 fused to mCherry-FKBP to the mitochondria and assessing delocalization of cDKA;1 fused with GFP. In this setup, KRP2-mCherry-FKBP could be anchored to the mitochondria, although the efficiency remained very low. We were able to find KRP2-positive mitochondria, in approximately 5% of cells imaged. In those events where mitochondrial targeting was observed however, this was not always correlated with CDKA;1 delocalization (compare the yellow arrowhead with the red arrowheads). These negative results point to limitations of delocalization-dependent visualization of protein-protein interactions due to putative problems concerning compartmentalization, protein stability, steric hindrance of the tags or competitive interaction strength. Scale bars = 20µm. This figure serves to counterbalance the positive results in Figure 3 by showing that not all interactions can be visualized with this system.

**Supplemental Table 1: List of primers used in this study.**

| Entry clone | Forward primer | Reverse primer | Reference |
|---|---|---|---|
| pDONR221-TagBFP2 | AttB1-GGGGACAAGTTTGTACAAAAAAGCAGGCTATGTCATCTAAGGGTGAAGAGCTTATCAAAGAGAAT | AttB2-GGGGACCACTTTGTACAAGAAAGCTGGGTCACCTCCGCCACCTCCACCTCCCAGTCCTGCGTA | (Pasin et al., 2014) |
| pDONR221-MITOTagBFP2 | AttB1-GGGGACAAGTTTGTACAAAAAAGCAGGCTCAATGAAGAGCTTCATTACAAGGAACAAGACAGCCATTTTGGCAACCGTTGCTGCTACAGGTACTGCCATCGGTGCCTACTATTATTACAACCAATTGCAACAGGATCCACCGGTCGCCACCATGTCATCTAAGGGTGAAGAGCTT | AttB2-GGGGACCACTTTGTACAAGAAAGCTGGGTACGCTAAGTCTTCCTCTGAAATCAA | (Robinson et al., 2010) |
| pDONR221-MITO | AttB1-GGGGACAAGTTTGTACAAAAAAGCAGGCTTTATGAAGAGCTTCATTACAAGGAACAAGACAGCCATTTTGGCAACCGTT | AttB2-GGGGACCACTTTGTACAAGAAAGCTGGGTAGGTGGCGACCGGTGGATCCTGTTGCAATT | (Robinson et al., 2010) |
| pDONR221-LHW | AttB1-GGGGACAAGTTTGTACAAAAAAGCAGGCTCGATGGGAGTTTTACTAAGAGAAGC | AttB2-GGGGACCACTTTGTACAAGAAAGCTGGGTGCATTGAACAGCCACCAGTAACCG | Bert De Rybel |
| pDONR221-SACL3 | AttB1-GGGGACAAGTTTGTACAAAAAAGCAGGCTCGATGCAGAACAATCAGTTTCCTC | AttB2-GGGGACCACTTTGTACAAGAAAGCTGGGTGAGATTGGTTTGAGAAATGTCC | Bert De Rybel |
| pDONR221-FKBP-mCherry | FWD-AttB1-FKBP-GGGGACAAGTTTGTACAAAAAAGCAGGCTCTATGGGAGTGCAGGTGGAAACCA | REV-FKBP-linker GGCCCCAGCGGCCGCAGCAGCACCAGCTTCCAGTTTTAGAAGCTCCACATCGAAG | (Robinson et al., 2010) |
| | FWD-linker-mCherry-GCTGGTGCTGCTGCGGCCGCTGGGGCCATGGTGAGCAAGGGCGAGGAGG | REV_AttB2-mCherry-GGGGACCACTTTGTACAAGAAAGCTGGGTACTTGTACAGCTCCTCCATGCCG | |
| pDONRP2RP3-mCherry-FKBP | FWD_AttB2-mCherry-GGGGACAGCTTTCTTGTACAAAGTGGCTATGGTGAGCAAGGGCGAGGAGG | REV_mCherry-linker-GGCCCCAGCGGCCGCAGCAGCACCAGCCTTGTACAGCTCCTCCATGCC | (Robinson et al., 2010) |
| | FWD_linker-rc-FKBP-GCTGGTGCTGCTGCGGCCGCTGGGGCCATGGGAGTGCAGGTGGAAACC | REV_AttB3-FKBP-GGGGACAACTTTGTATAATAAAGTTGTTTATTCCAGTTTTAGAAGCTCCACATCG | |
| pDONRP2RP3-FRB* | AttB2r-GGGGACAGCTTTCTTGTACAAAGTGGGGATCCTCTGGCATGAGATGTGG | AttB3-GGGGACAACTTTGTATAATAAAGTTGTTACTTTGAGATTCGTCGGAACACATG | (Chang et al., 2013, 2017) |
| pENTR5'TOPO-UBQ10NLS | FWD_pUBQ10-CGACGAGTCAGTAATAAACGGCG | REV_NLS-CCACCTCCGCCACCTCCTCCAACCTTTCTCTTC | (Mehlmer et al., 2012) |

**Supplemental Table 2: List of constructs used in this study.**

| Entry clones | | |
|---|---|---|
| **Clone name** | **Accession number** | **Reference** |
| pDONR207 MAP65-1 | At5G55230 | (Van Damme et al., 2004) |
| pDONR207-CLC1 | At2G20760 | (Van Damme et al., 2011) |
| pDONR207-RCN1/PP2AA1 | At1G25490 | (Spinner et al., 2013) |
| pDONR207-T22.1/TPLATE | At3G01780 | (Van Damme et al., 2004) |
| pDONR207-EB1a | At3G47690 | (Van Damme et al., 2004) |
| pDONR221-BIN2 | At4G18710 | (Houbaert et al., 2018) |
| pDONR221-CDKA;1 | At3G48750 | (Boruc et al., 2010) |
| pDONR221-KRP2 | At3G50630 | (Boruc et al., 2010) |
| pDONR221-LHW | At2G27230 | Bert De Rybel |
| pDONR221-MITO | C8ZGB0 | this work, (Robinson et al., 2010) |
| pDONR221-MITOTagBFP2 | C8ZGB0 | this work |
| pDONR221-MYRI | At5G19360 | (Kirik et al., 2012; Myers et al., 2009; Podell and Gribskov, 2004) |
| pDONR221-SACL3 | At1G29950 | Bert De Rybel |
| pDONR221-TagBFP2 | this work | (Pasin et al., 2014) |
| pDONR221-TASH3 | At2G07360 | (Gadeyne et al., 2014) |
| pDONR221-FKBP-mCherry | this work | this work, (Mylle et al., 2013; Robinson et al., 2010) |
| pDONRP2RP3-mCherry-FKBP | this work | this work, (Mylle et al., 2013; Robinson et al., 2010) |
| pDONRP2RP3-FRB* | this work | this work, (Chang et al., 2013, 2017) |
| pDONRP2RP3-Sac1 | YKL212W | (Doumane and Caillaud, 2020) |
| pDONRP2RP3-Sac1-dead | YKL212W | (Doumane and Caillaud, 2020) |
| pENTR 5' TOPO pUBQ10-NLS | this work | this work, (Mehlmer et al., 2012) |
| **Expression clones** | | |
| **Clone name** | **Accession number** | **Reference** |
| p35S:BES1-GFP | At1G19350 | Eugenia Russinova |
| pB7m34GW:p35S:CDKA;1-mCherry-FKBP | At3G48750 | this work |
| pB7m34GW:p35S:CLC1-mCherry-FKBP | At2G20760 | this work |
| pB7m34GW:p35S:LHW-mCherry-FKBP | At2G27230 | this work |
| pB7m34GW:p35S:MAP-65-1-mCherry-FKBP | At5G55230 | this work |
| pB7m34GW:p35S:MYRI-TagBFP2-FRB* | At5G19360 | this work |
| pB7m34GW:p35S:RCN1/PP2AA1-mCherry-FKBP | At1G25490 | this work |
| pB7m34GW:p35S:SACL3-eGFP | At1G29950 | this work |
| pB7m34GW:p35S:TASH3-TagBFP2 | At2G07360 | this work |
| pB7m34GW:p35S:TPLATE-mCherry-FKBP | At3G01780 | this work |
| pB7m34GW:pUBQ:GFP-AUXILIN-LIKE1 | At4G12780 | (Adamowski et al., 2018) |
| pB7m43GW:p35S:EB1A-TagBFP-FRB* | At3G47690 | this work |
| pGWB6:GFP-FASS/TON2 | At5G18580 | (Spinner et al., 2013) |

| | | |
|---|---|---|
| pK7FWG2:p35S:LOLITA-GFP | At1G15370 | (Gadeyne et al., 2014) |
| pK7FWG2:p35S:TML-GFP | At5G57460 | (Gadeyne et al., 2014) |
| pK7FWG2:p35S:MAP65-1-GFP | At5G55230 | (Van Damme et al., 2004) |
| pK7FWG2:p35s:CDKA;1-GFP | At3G48750 | (Boruc et al., 2010) |
| pK7WG2:p35S:KRP2-GFP | At3G50630 | (Boruc et al., 2010) |
| pB7m43GW:p35S:KRP2-mCherry-FKBP | At3G50630 | this work |
| pK7WG2:eGFP | this work | (Karimi et al., 2002) |
| UBQ10prom:mCitrine-P4M | DQ845395 | (Doumane and Caillaud, 2020; Simon et al., 2016) |
| pB7m34GW:p35S:FKBP-mCherry-Sac1 | YKL212W | this work |
| pB7m34GW:p35S:FKBP-mCherry-Sac1 | YKL212W | this work |
