## Supplemental File 1 for "Visualizing protein-protein interactions in plants by rapamycin-dependent delocalization"

### Manual of the quantification scripts

This supplemental document describes procedures for proper data processing for the rapamycin-dependent delocalization assay. The document is divided into two sections, A and B, which correspond to either mitochondrial (MitTally) or nuclear (NucTally) subcellular delocalization. Prior to the analysis user should validate the pictures for the saturation. Pictures that contain saturated particles/nuclei, should not be analyzed. Both scripts are written in groovy programing language and are compatible with Fiji/ImageJ.

A) MitTally (Mitochondria Tally marks) – the script is automatically calculating intensity ratios of a particular channel 'particle' (mitochondrion) area versus the cell area within the user's pre-defined region of interest (ROI). The cell area excludes particle areas from the calculation. Particle areas are generated on the basis of the 'Mito channel' via automatic mitochondria recognition. Analysis should be performed as follows:

1. Open Fiji/ImageJ.
2. Open the ROI Manager **Analyze>Tools>ROI Manager**.
3. Load the picture for the quantification.
4. Outline the cell for quantification as accurately as possible with the **Freehand selections tool** or the **Polygon selection tool**, avoiding the nuclear area (see Screenshot 1). Make sure to avoid the saturated PM of neighboring cells.
5. **Add** the ROI to the ROI Manager.
6. Select the ROI and in ROI Manager and click **More>Save**
7. Save the ROI in the same folder as the source picture under the same name, with *.roi extension (see Screenshot 2).
8. Perform ROI creation for all of the pictures you want to quantify.
9. Load the script into Fiji/ImageJ and from the script Language menu, choose Groovy.
10. In the script window, click 'Run' (see Screenshot 3).
11. Choose the settings in the pop-up window (see Screenshot 4):
    - Input directory – select the location of the folder containing the pictures for analysis (see Note 1),
    - Mito channel – select the number of the channel that contains the mitochondrial construct expression (see Note 2),

- Red channel/Green channel/Blue channel – select the number of the channel corresponding to the fluorophore or 'None' (see Note 3),
- Minimum particle size/Maximum particle size – corresponds to the size of particle in pixels (see Note 4)
- Maxima tolerance – value which will determine, the level of separation of particles via watershed function (see Screenshot 5 and Note 5)
- Threshold Method for mitochondria – select the thresholding method for mitochondria recognition (see Note 6).

12. Press 'Ok'.
13. The script automatically recognizes the mitochondria in the corresponding 'Mito Channel' and generates a ROI for each of the found particles.
14. A pop-up window notifies when the calculations are finished (see Screenshot 7).
15. The script generates a result file for each of the analyzed pictures as well as one summarizing results file for all of the analyzed pictures and session settings summary file. Also, *.zip files containing ROIs of all identified mitochondria ('image name'_mito.zip) and cell ROI excluding mitochondria ('image name'_cell_minus_mito.zip) are generated. The mitochondria ROIs can be drag and dropped as *.zip file directly to ImageJ/Fiji to visualize all particles at once. If during opening the Excel file the error windows pop-up, saying 'The file format and extension of '*file name*' don't match. The file could be corrupted or unsafe. Unless you trust its source, do not open it. Do you want to open it anyway?', click 'Yes'. After opening the file, it can be saved as new one in other Microsoft Excel formats.
16. In the individual excel file ('*image name*'-results.xls) the following data are generated:

    FileName, Particle Id, Particle Area, Cell Area, Cell Red Mean, Cell Red Median, Cell Green Mean, Cell Green Median, Cell Blue Mean, Cell Blue Median, Mean Red Particle, Median Red Particle, Mean Green Particle, Median Green Particle, Mean Blue Particle, Median Blue Particle, Mean Red Ratio (P/C), Median Red Ratio (P/C), Mean Green Ratio (P/C), Median Green Ratio (P/C), Mean Blue Ratio (P/C), Median Blue Ratio (P/C). 'Particle' and 'cell' areas are given in pixels. Cells and particles *color* 'mean' and 'median' are corresponding to the area mean or median intensity values.

'P/C' is a ratio of certain particle area divided by cell area (subtracted of all particle areas). The file can be opened in Microsoft Excel.

17. The excel summary result file (result-Mito-summary-'*folder name*'-'*used threshold method name*'.xls) provides the following data:
    FileName, # of Particles, Red P/C mean, Green P/C mean, Blue/PC mean, Red P/C median, Green P/C median, Blue/PC median, # of Particles Filtered, Red P/C mean Filtered, Green P/C mean Filtered, Blue/PC mean Filtered, Red P/C median Filtered, Green P/C median Filtered, Blue/PC median Filtered. '# of Particles' corresponds to the total number of mitochondria identified by script in each image. Mean/median values of all P/C ratios of particular *color* in each picture are average values calculated on the basis of mean or median (correspondingly) particles and cell areas intensities. 'Filtered' corresponds to results calculated for all data points excluding the outliers. Outliers are defined on the basis of the single-step interquartile range (IQR), and are identified per picture on the basis of particles mean or median intensity value.
18. The text summary result file (result-Mito-summary-'*folder name*'-'*used threshold method name*'.txt) provides information about the session settings chosen by the user.
19. We recommend to check the script-generated particles areas ('image name'_mito.zip) via drag-drop into Fiji/ImageJ to ensure, that user-selected settings were suitable for the pictures analyzed in the session.
20. If the particle detection in a few of the pictures out of all analyzed is not done properly, we recommend splitting up the database and perform the analysis separately.

B) NucTally (Nuclear Tally marks) – the script is automatically calculating intensity ratios of a particular channel 'nucleus' area versus the cell area within user's pre-defined region of interest (ROI). The cells areas exclude nuclei areas. Nuclei areas are generated on the basis of the 'Nucleus channel' automatically or can be pre-defined by user (see Note 7). Analysis should be performed as follows:

1. Open Fiji/ImageJ.
2. Open the ROI Manager **Analyze>Tools>ROI Manager**.
3. Load the picture for the quantification.
4. Outline the cell for quantification as accurately as possible with the **Freehand selections tool** or the **Polygon selection tool** (see Screenshot 8). Make sure to avoid the saturated PM of neighboring cells. Make sure to outline only one cell, otherwise all nuclei present in the ROI will be identified and analyzed in the later steps together.
5. **Add** the ROI to the ROI Manager.
6. Select the ROI in the ROI Manager and click **More>Save**
7. Save the ROI in the same folder as the source picture under the same name, with *.roi extension (see Screenshot 2).
8. (Optional) Outline the nucleus and **Add** the ROI to the ROI Manager. Save the ROI in the same folder as the source picture under the same name plus '_nucleus', with *.roi extension (see Screenshot 9),

   OR

   (Optional) Outline the nucleolus and **Add** the ROI to the ROI Manager.

   In the Roi Manager, select both nuclear and nucleolar ROIs. Click ROI menu **More>XOR>Add** to create a ROI of the nucleus excluding the nucleolus (see Screenshot 10). Save the ROI in the same folder as the source picture under the same name plus '_nucleus', with *.roi extension (see Note 8).
9. Perform ROI creations for all pictures you want to quantify.
10. Load the script into Fiji/ImageJ and from the script Language menu, choose Groovy.
11. In the script window, click 'Run' (see Screenshot 11).
12. Choose the settings in the pop-up window (see Screenshot 12).
    - Select a directory – select the location of the folder containing the pictures for analysis (see Note 1),
    - Minimum Nucleus ROI area/ Minimum Nucleus ROI area – corresponds to the size of nuclei in pixels (see Note 9)
    - Nucleus channel – select number of channel with nuclear construct expression, or ROI if nuclei ROIs were user-generated (see Note 7 and Note 8),

- Red channel/Green channel/Blue channel – select the number of the channel corresponding to the fluorophore or 'None' (see Note 3),
- Nucleus Threshold Method – select the thresholding method for nuclei recognition,
- Use American Decimal – check or uncheck, to use either '.' or ',' as decimal.

13. Press 'Ok'.
14. The script recognizes nuclei in the corresponding 'Nucleus Channel' and generates a ROI for each or is using user pre-defined nuclei ROIs.
15. Prior to calculations, the script is shrinking the nuclei area from the outside (function *Erode*, applied twice, one pixel each time) in order to avoid including the nuclear envelope in the analysis.
16. A pop-up window notifies when the calculations are finished (see Screenshot 13).
17. The script generates one summary result file for all of the analyzed pictures, two summaries results files for all of the analyzed pictures excluding outliers and session settings summary file. If during opening the Excel file the error windows pop-up, saying 'The file format and extension of '*file name*' don't match. The file could be corrupted or unsafe. Unless you trust its source, do not open it. Do you want to open it anyway?', click 'Yes'. After opening the file, it can be saved as new one in other Microsoft Excel formats.
18. In the results summary excel file (result-Nucleus-summary-'*folder name*' -'*used threshold method name*'.xls) the following data are generated:

    File Name, Cell Area, Nucleus Area, Mean Intensity Cell Red, Mean Intensity Nucleus Red, Mean Intensity Cell Green, Mean Intensity Nucleus Green, Mean Intensity Cell Blue, Mean Intensity Nucleus Blue, Red Mean Intensity Ratio (N/C), Green Mean Intensity Ratio (N/C), Blue Mean Intensity Ratio (N/C), Median Intensity Cell Red, Median Intensity Nucleus Red, Median Intensity Cell Green, Median Intensity Nucleus Green, Median Intensity Cell Blue, Median Intensity Nucleus Blue, Red Median Intensity Ratio (N/C), Green Median Intensity Ratio (N/C), Blue Median Intensity Ratio (N/C). Cells and nuclei *color* mean and median are corresponding to the area mean or median intensity values. N/C is a ratio of certain nucleus area divided by cell area (subtracted of nucleus area).

19. In the results summaries Excel files (result-Nucleus-Mean-Without-Outlier -'*folder name*' -'*used threshold method name'*.xls and result-Nucleus-Median-Without-Outlier -'*folder name*' -'*used threshold method name'*.xls) the following data are generated: File Name, Red N/C mean (or)median Filtered, Green N/C mean (or) median Filtered, Blue N/C mean (or) median Filtered. 'N/C' corresponds to ratios of particular *color* in each picture calculated on the basis of mean or median (correspondingly) nuclei and cell areas intensities. In these files there are presented pictures that were not identified as outliers. Outliers are defined on the basis of the single-step interquartile range (IQR), and are identified per dataset on the basis of N/C ratios calculated from mean or median intensity values.
20. The text summary result file (result-Nucleus-summary-'*folder name*' -'*used threshold method name*'.txt) provides information about the session settings chosen by the user.

Note 1. The location cannot contain any subfolders.

Note 2. All pictures from a single directory have to have the same number of channels. We also recommend either performing sequential imaging keeping the same channel order (e.g. 488nm Green, 561nm Red, 405nm Blue), or rearranging them via *I*mageJ>Image>Color>Arrange Channels (can be performed only on pictures with multiple channels: Channels tool Ctrl+Shift+Z, change Composite to Color).

Note 3. If 'None' is selected, quantification will not be performed for that channel.

Note 4. Particles, which do not fit in the user's selected range, will not be recognized by the script. We recommend to start by using the default settings.

Note 5. Maxima tolerance is a setting, which uses the Watershed Fiji/ImageJ plug-in to determine how much the found particle will be segmented. It is useful to identify clustered particles as individual ones.  The higher the value the less the particle will be segmented. In addition, the more homogenous particle's intensity is, the less segmented it will be, regardless of Maxima Tolerance settings (see Screenshot 5). We recommend to use a default value of 2300.

Note 6. CRUCIAL STEP. Selection of the proper thresholding method depends on multiple factors, of which the most important is the signal to noise ratio of the mitochondria and the background. Therefore, we do recommend testing thresholding methods prior the selection in script on a few exemplary pictures from the same experiment. To do so, open the picture in Fiji/ImageJ, load the picture with its cell ROI and select the channel with the mitochondria. In the threshold menu (Image>Adjust>Threshold) apply different threshold methods. Select the one, which guarantees good signal to noise resolution while accommodating the maximal number of mitochondria of proper size (see Screenshot 6).

Note 7. We recommend automatic nuclear recognition only with those pictures, where the nuclear construct expression is strong and easily distinguishable from the background.

Note 8. We recommend user-creation of nuclei area, if the nuclear construct is not highly expressed and hardly distinguishable from the background. We recommend subtracting the nucleolar area from nucleus if a high accumulation of signal is observed in the nucleolus, and if this leads to saturation of the pixels in that area.

Note 9. If selected values do not correspond with the actual nuclei sizes, analysis will not be performed. We recommend using the default settings.

Screenshot 1. Exemplary proper cell outlining for MitTally analysis. Note, the nucleus should not be included in the ROI, if protein(s) of interests exhibit strong nuclear expression.

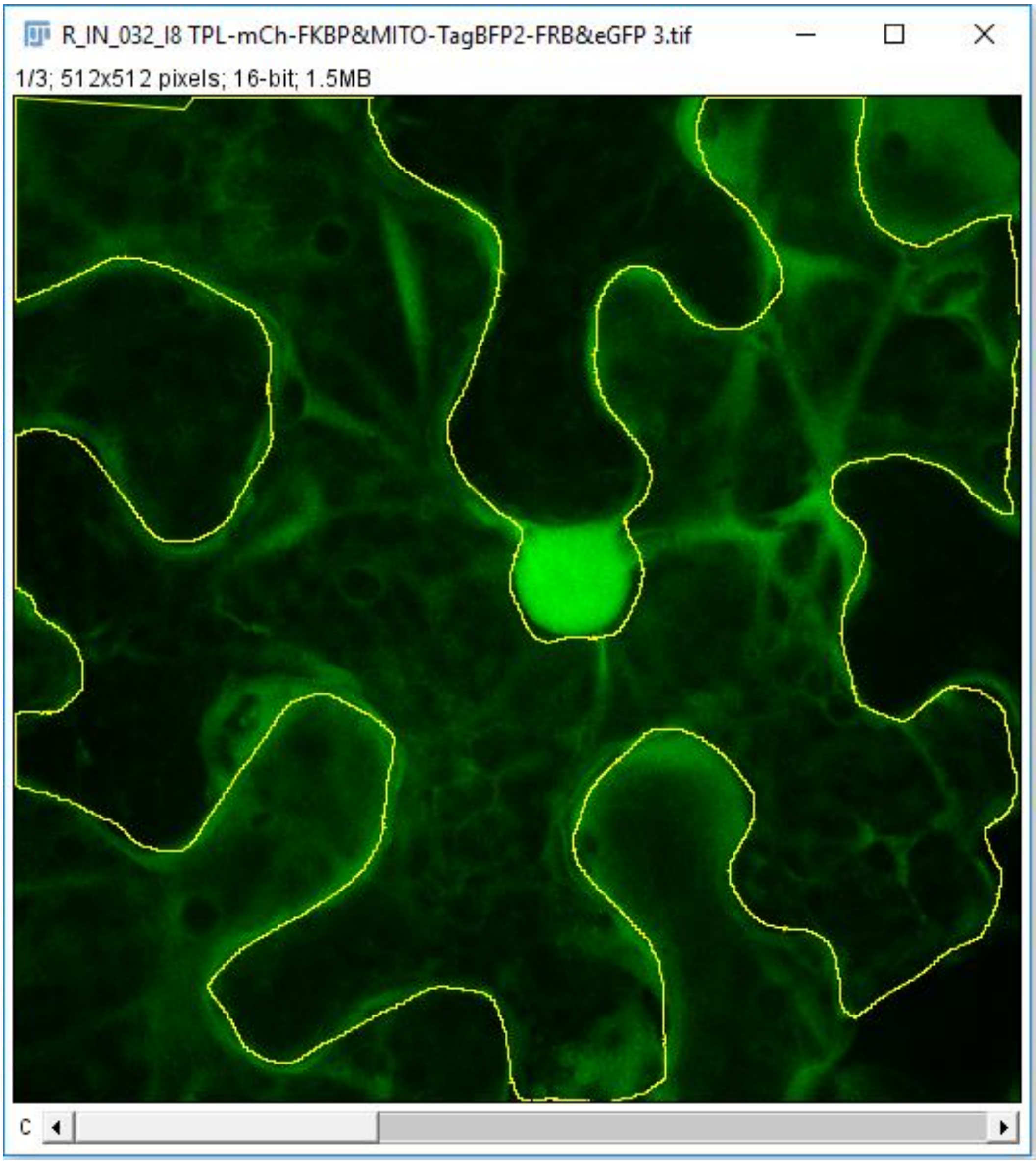

Screenshot 2. List of computed pictures with paired ROI files. Remark, ROI files should hold the same name as the corresponding picture.

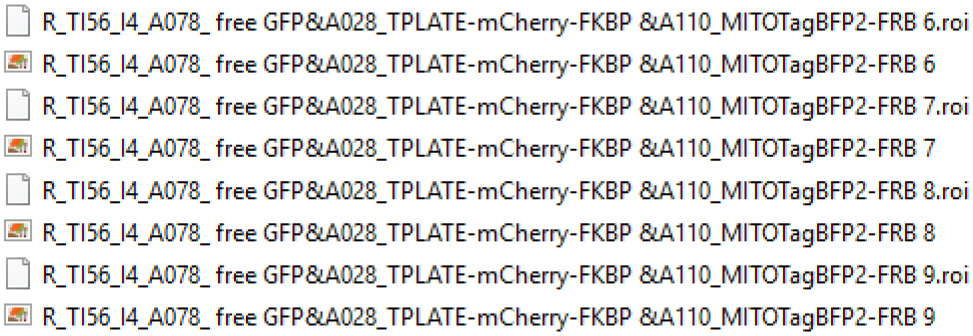


Screenshot 3. MitTally script loaded in Fiji/ImageJ.

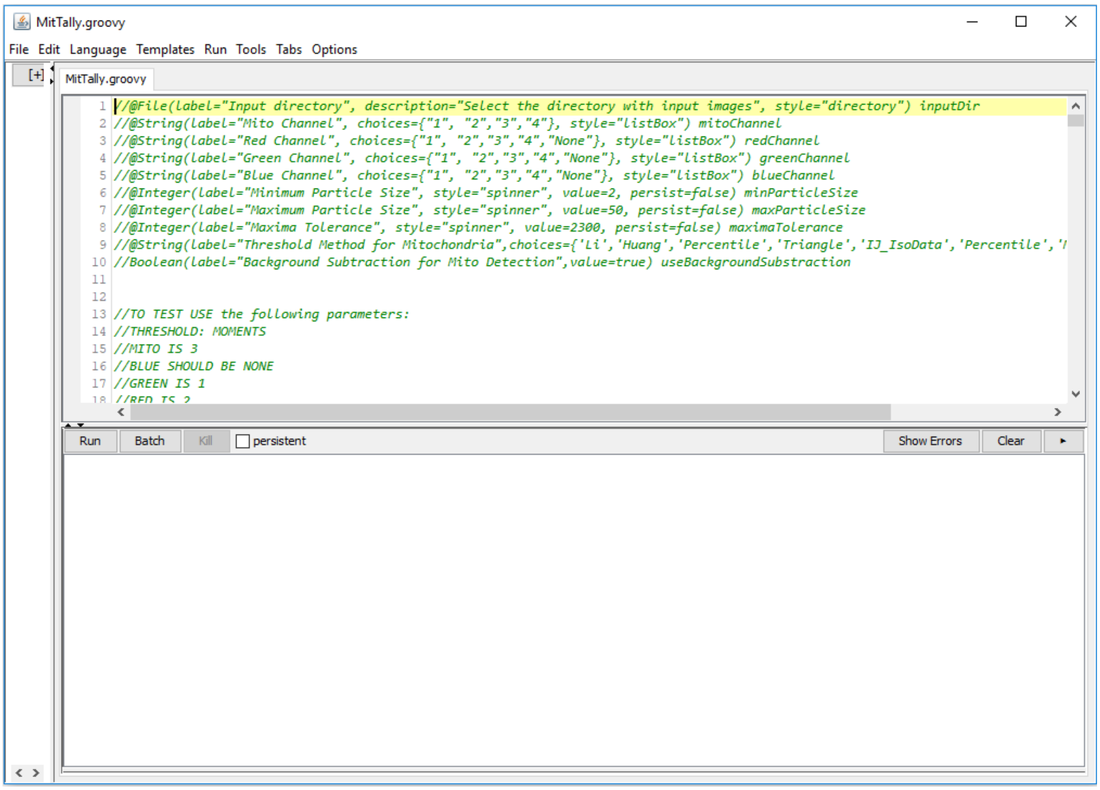

Screenshot 4. MitTally settings window.

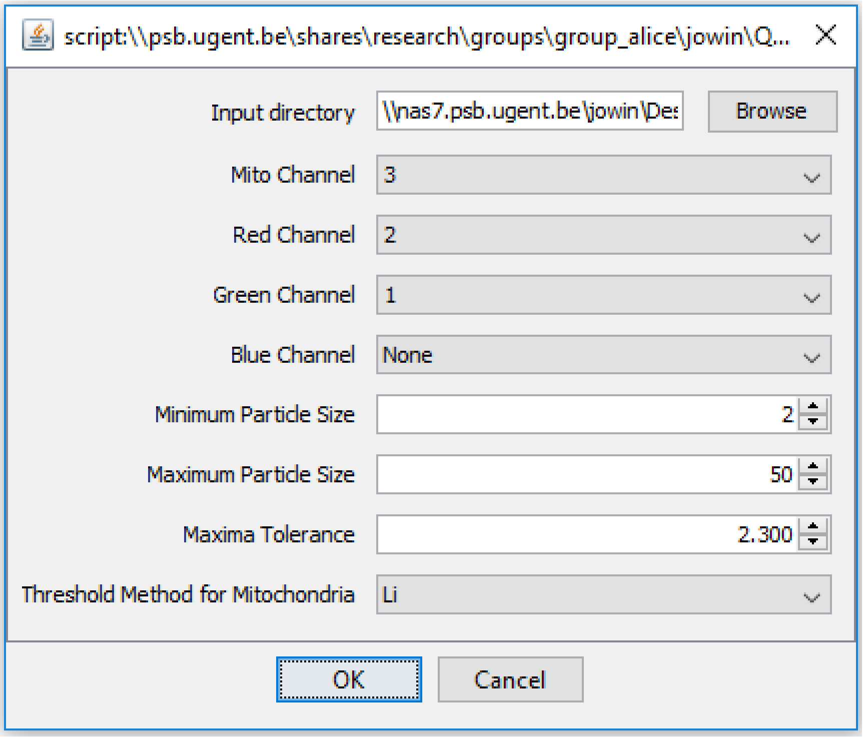

Screenshot 5. Comparison of segmentation with a use of different Maxima Tolerance value. Remark, the more the heterogeneous the particle’s intensity is in different area, the more segmented it will be (A). Contrary, the more homogenous intensity, the less segmentation will be computed regardless of Maxima Tolerance settings (B).

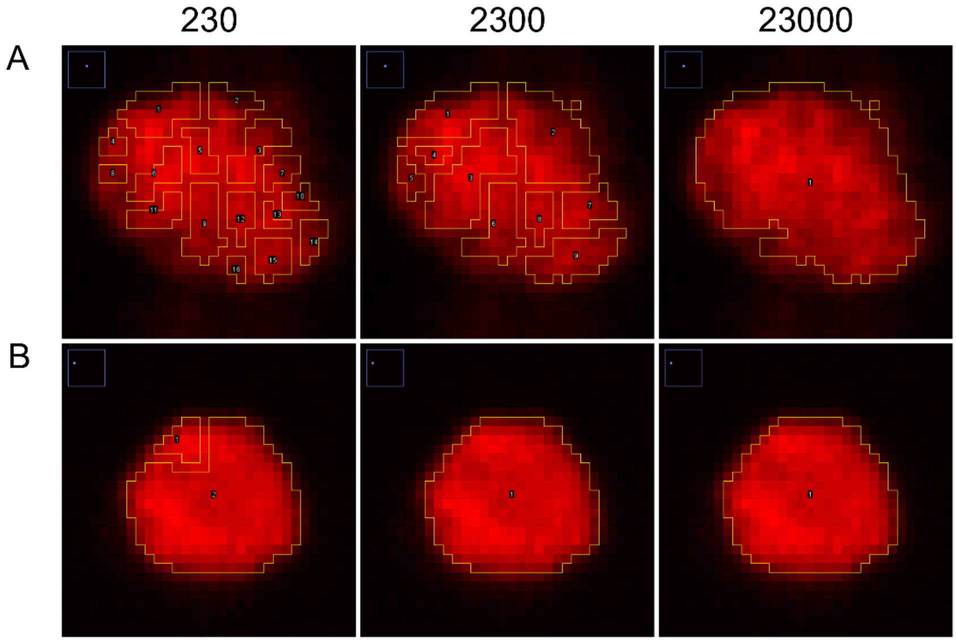

Screenshot 6. Example of the proper selection of thresholding method. Upper left – background displays too high noise. Upper right – to few mitochondria are recognized. Bottom left – satisfying number of mitochondria are recognized, but their ROI area is too extensive. Bottom right - satisfying number of mitochondria are recognized, and their ROI areas correspond with their actual size. Change only the threshold method (red arrowheads), do not change the minimum and maximum threshold bars.

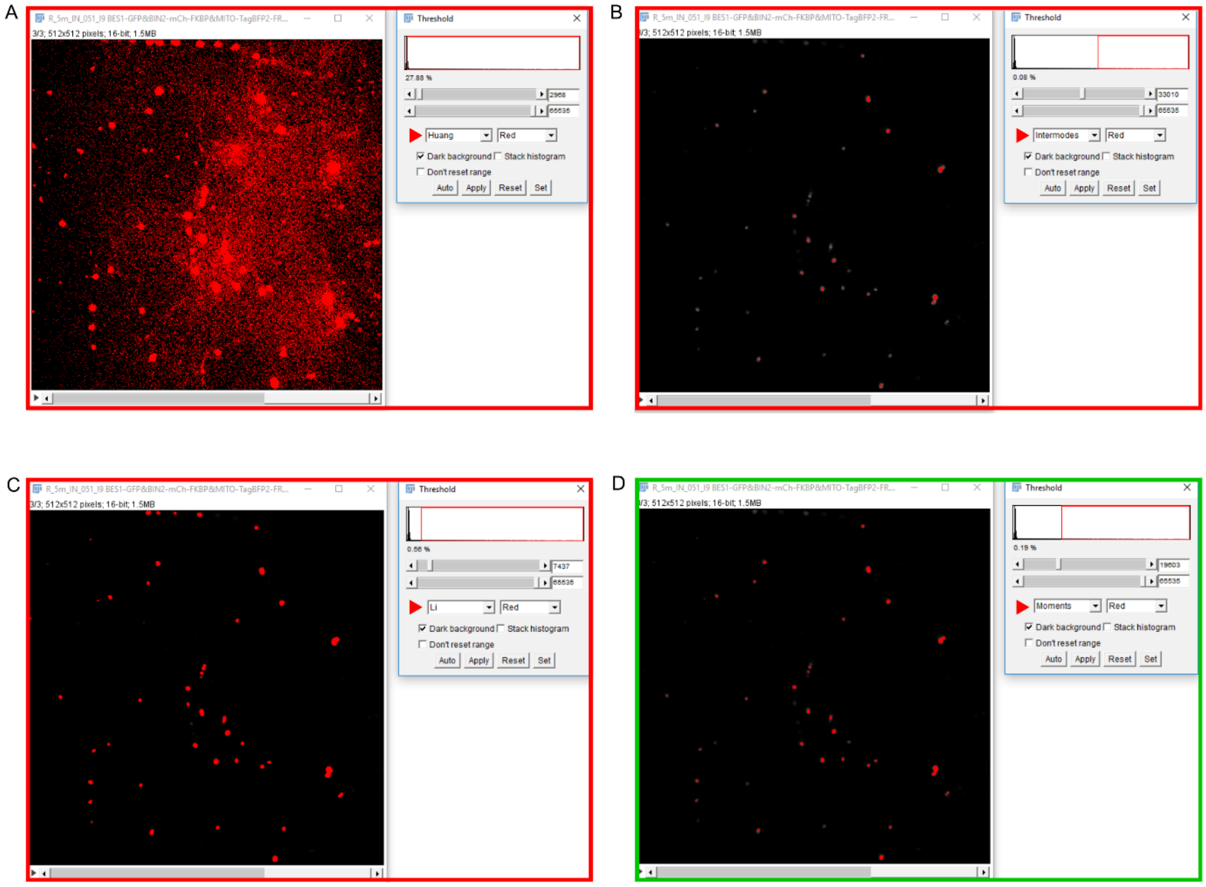

Screenshot 7. Pop-up window communicating completion of MitTally analysis.

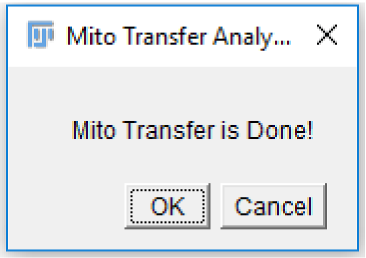


Screenshot 8. Exemplary proper cell outlining for NucTally analysis.

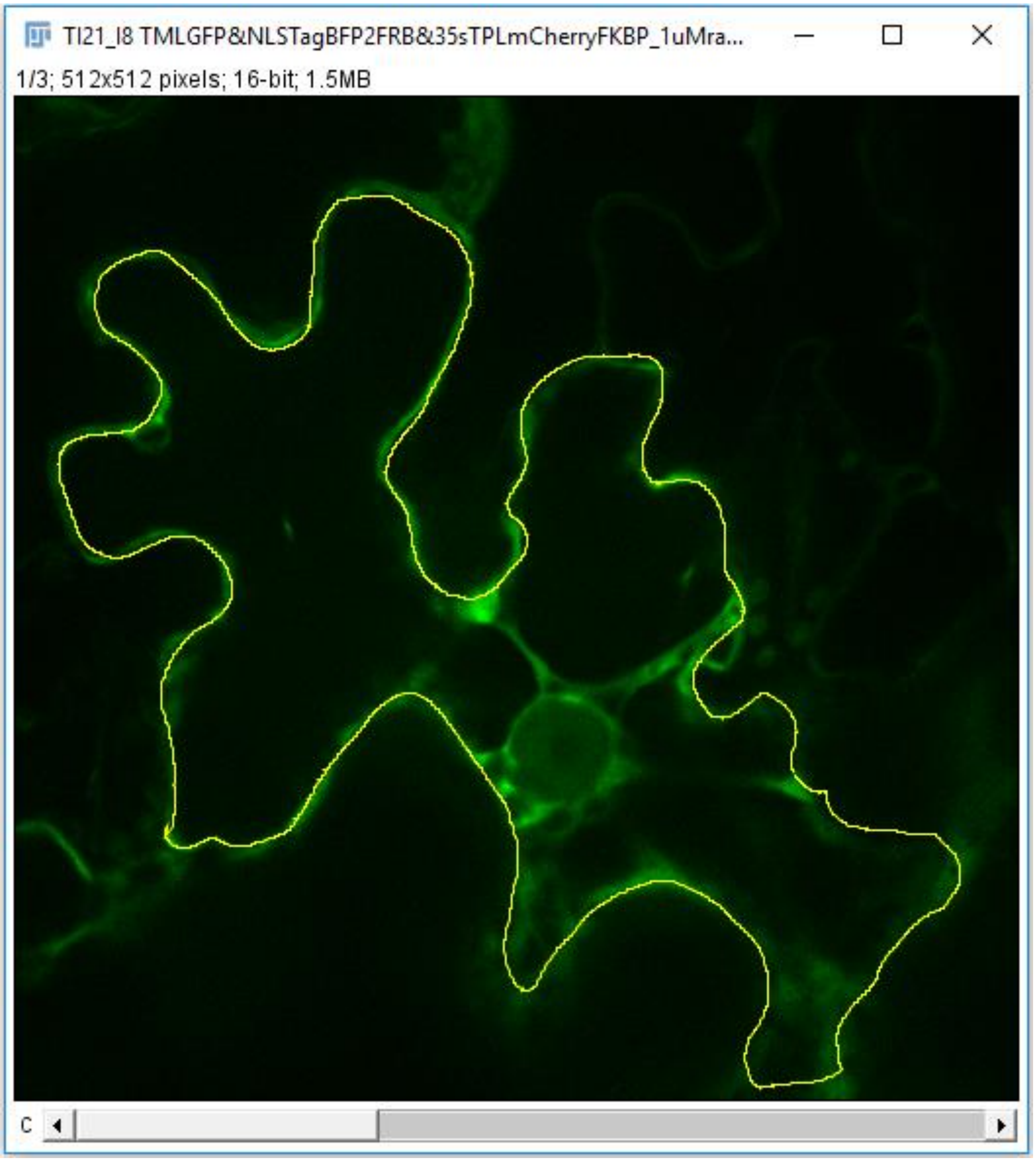

Screenshot 9. List of computed pictures with paired ROI files. Remark, ROI files should hold the same name as the corresponding picture.

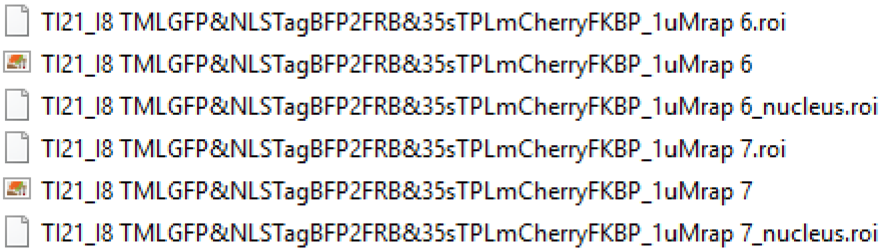


Screenshot 10. Steps of creating ROI of nucleus excluding the nucleolus. Upper left – outline nucleus and Add ROI to the ROI Manager. Upper right – outline nucleolus and Add ROI to the ROI Manager. Bottom left - select nucleus and nucleolus ROIs, use More>XOR function. Bottom right – delete nucleus and nucleolus ROIs, More>Save XOR ROI.

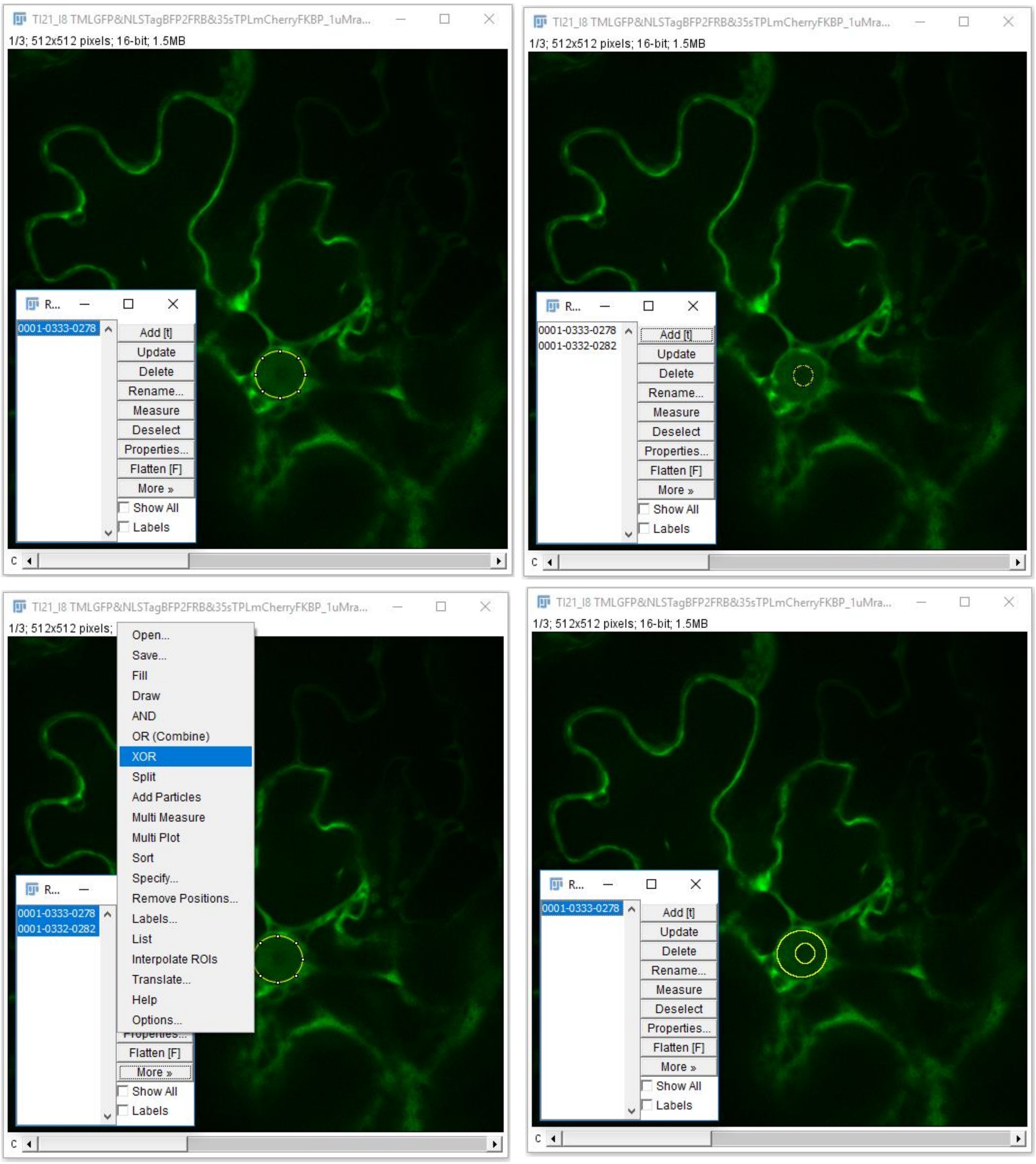

Screenshot 11. NucTally script loaded in Fiji/ImageJ.

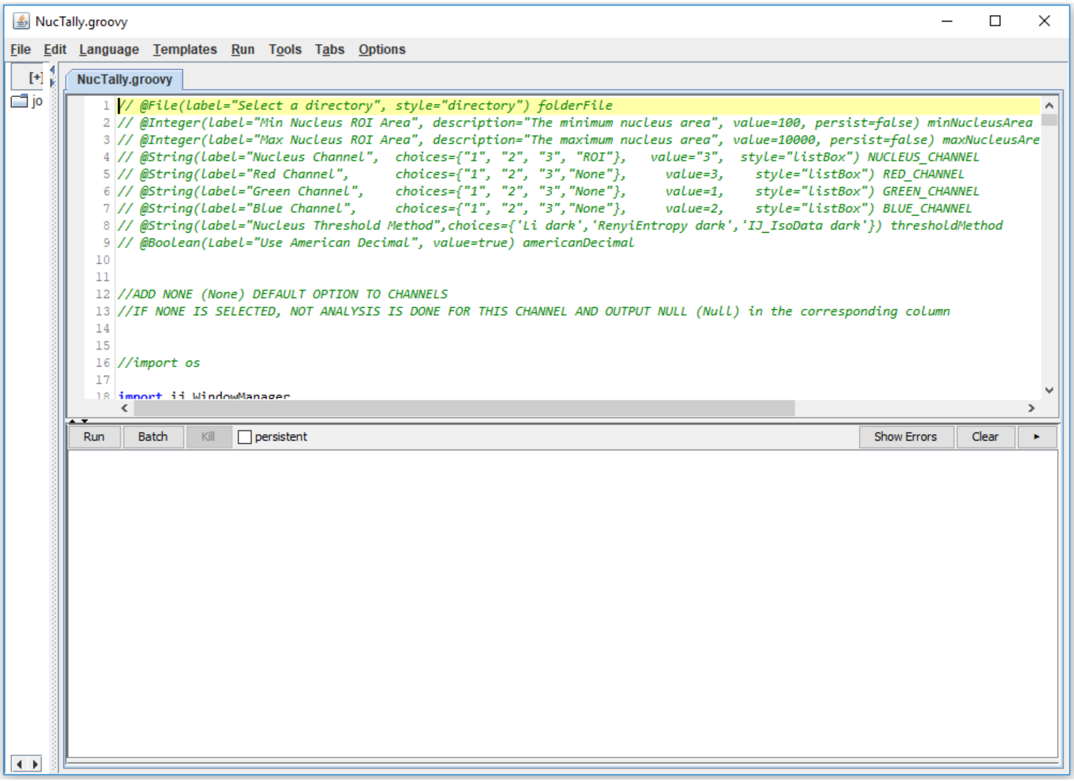


Screenshot 12. NucTally settings window.

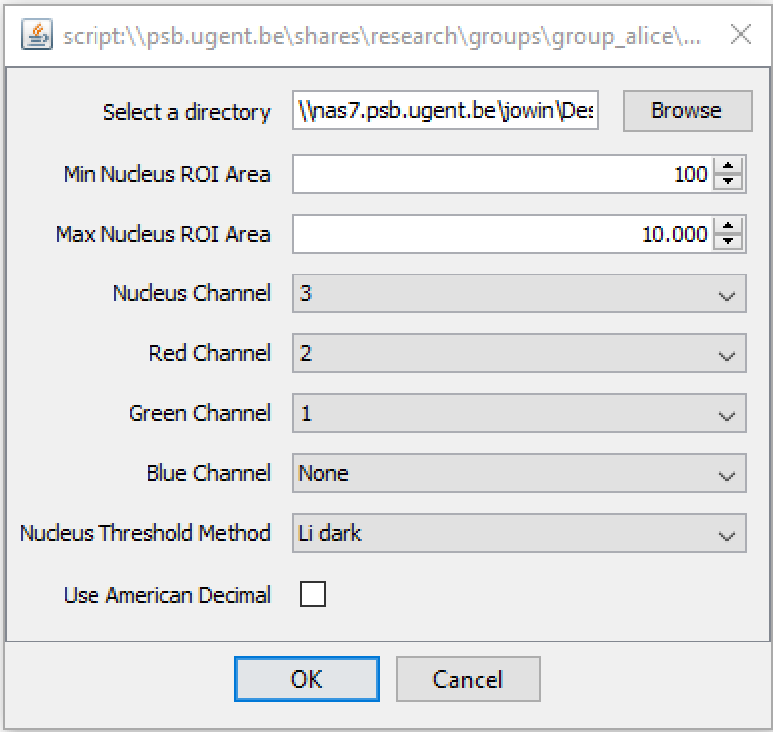

Screenshot 13. Pop-up window communicating completion of NucTally analysis.

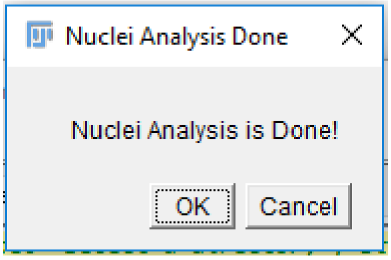
