## Supplemental File 4 for "Visualizing protein-protein interactions in plants by rapamycin-dependent delocalization"

Supplemental Data. Winkler et al. (2021). Visualizing protein-protein interactions in plants by rapamycin-dependent delocalization. Plant Cell.

**Supplemental File 4. Statistical analysis**

**Figure 2C**

Std. Error = Standard Error, SAC= FKBP-mCherry-Sac1, SAC_D = FKBP-mCherry-Sac1-dead, _UNTR = untreated, _RAP = rapamycin treated

| | Estimate | Std. Error | t value | Pr(>\|t\|) |
|---|---|---|---|---|
| SAC_D_UNTR - SAC_D_RAP | -0.17733 | 0.08742 | -2.028 | 0.150 |
| SAC_RAP - SAC_D_RAP | 25.41035 | 3.46299 | 7.338 | <0.001 *** |
| SAC_UNTR - SAC_D_RAP | 0.12729 | 0.16919 | 0.752 | 0.848 |
| SAC_RAP - SAC_D_UNTR | 25.58768 | 3.46195 | 7.391 | <0.001 *** |
| SAC_UNTR - SAC_D_UNTR | 0.30462 | 0.14645 | 2.080 | 0.135 |
| SAC_UNTR - SAC_RAP | -25.28306 | 3.46498 | -7.297 | <0.001 *** |

---

Signif. codes: 0 '***' 0.001 '**' 0.01 '*' 0.05 '.' 0.1 ' ' 1

(Adjusted p values reported -- single-step method)

**Figure 3Ai**

Std. Error = Standard Error, aTPLATE_R = TPLATE-mCherry-FKBP rapamycin treated, aTPLATE = TPLATE-mCherry-FKBP untreated, fGFP = free GFP untreated, fGFP_R = free GFP rapamycin treated

| | Estimate | Std. Error | t value | Pr(>\|t\|) |
|---|---|---|---|---|
| aTPLATE_R - aTPLATE | 1.96983 | 0.46462 | 4.240 | <0.001 *** |
| fGFP - aTPLATE | -0.08867 | 0.12371 | -0.717 | 0.878 |
| fGFP_R - aTPLATE | -0.29204 | 0.11688 | -2.499 | 0.060 . |
| fGFP - aTPLATE_R | -2.05850 | 0.46338 | -4.442 | <0.001 *** |
| fGFP_R - aTPLATE_R | -2.26187 | 0.46161 | -4.900 | <0.001 *** |
| fGFP_R - fGFP | -0.20337 | 0.11188 | -1.818 | 0.248 |

---

Signif. codes: 0 '***' 0.001 '**' 0.01 '*' 0.05 '.' 0.1 ' ' 1

(Adjusted p values reported -- single-step method)

**Figure 3Bi**

Std. Error = Standard Error, LHW_R = LHW-mCherry-FKBP rapamycin treated, LHW = LHW-mCherry-FKBP untreated, SACL3 = SACL3-GFP untreated, SACL3_R = SACL3-GFP rapamycin treated

| | Estimate | Std. Error | t value | Pr(>\|t\|) |
|---|---|---|---|---|
| LHW_R - LHW | 2.39354 | 0.27842 | 8.597 | <0.001 *** |
| SACL3 - LHW | 0.10158 | 0.06531 | 1.555 | 0.369 |
| SACL3_R - LHW | 2.62088 | 0.32124 | 8.159 | <0.001 *** |
| SACL3 - LHW_R | -2.29196 | 0.28061 | -8.168 | <0.001 *** |
| SACL3_R - LHW_R | 0.22733 | 0.42151 | 0.539 | 0.940 |
| SACL3_R - SACL3 | 2.51930 | 0.32314 | 7.796 | <0.001 *** |

---

Signif. codes: 0 ‘***’ 0.001 ‘**’ 0.01 ‘*’ 0.05 ‘.’ 0.1 ‘ ’ 1

(Adjusted p values reported -- single-step method)

**Figure 3Ci**

Std. Error = Standard Error, aBIN2 = BIN2-mCherry-FKBP rapamycin treated, aBIN2 = BIN2-mCherry-FKBP untreated, BES1 = BES1-GFP untreated, BES1_R = BES1-GFP rapamycin treated

| | Estimate | Std. Error | t value | Pr(>\|t\|) |
|---|---|---|---|---|
| aBIN2_R - aBIN2 | 3.892619 | 1.170996 | 3.324 | 0.0115 * |
| BES1 - aBIN2 | 0.007143 | 0.094944 | 0.075 | 0.9998 |
| BES1_R - aBIN2 | 2.029286 | 0.682813 | 2.972 | 0.0243 * |
| BES1 - aBIN2_R | -3.885476 | 1.169231 | -3.323 | 0.0116 * |
| BES1_R - aBIN2_R | -1.863333 | 1.350674 | -1.380 | 0.4720 |
| BES1_R - BES1 | 2.022143 | 0.679783 | 2.975 | 0.0244 * |

---

Signif. codes: 0 ‘***’ 0.001 ‘**’ 0.01 ‘*’ 0.05 ‘.’ 0.1 ‘ ’ 1

(Adjusted p values reported -- single-step method)

**Figure 3Di**

Std. Error = Standard Error, aTPLATE = TPLATE-mCherry-FKBP rapamycin treated, aTPLATE = TPLATE-mCherry-FKBP untreated, TML = TML-GFP untreated, TML_R = TML-GFP rapamycin treated

| | Estimate | Std. Error | t value | Pr(>\|t\|) |
|---|---|---|---|---|
| aTPLATE_R - aTPLATE | 3.11416 | 0.39712 | 7.842 | <0.001 *** |
| TML - aTPLATE | -0.01294 | 0.08938 | -0.145 | 0.9987 |
| TML_R - aTPLATE | 1.98216 | 0.30372 | 6.526 | <0.001 *** |
| TML - aTPLATE_R | -3.12710 | 0.39577 | -7.901 | <0.001 *** |
| TML_R - aTPLATE_R | -1.13200 | 0.49081 | -2.306 | 0.0903 . |
| TML_R - TML | 1.99510 | 0.30195 | 6.607 | <0.001 *** |

---

Signif. codes: 0 ‘***’ 0.001 ‘**’ 0.01 ‘*’ 0.05 ‘.’ 0.1 ‘ ’ 1

(Adjusted p values reported -- single-step method)

**Figure 4Ai**

Std. Error = Standard Error, aTPLATE = TPLATE-mCherry-FKBP rapamycin treated, aTPLATE = TPLATE-mCherry-FKBP untreated, TML = TML-GFP untreated, TML_R = TML-GFP rapamycin treated

| | Estimate | Std. Error | t value | Pr(>\|t\|) |
|---|---|---|---|---|
| aTPLATE_R - aTPLATE | 1.88044 | 0.21826 | 8.616 | < 0.001 *** |
| TML - aTPLATE | -0.08071 | 0.15639 | -0.516 | 0.95257 |
| TML_R - aTPLATE | 0.82188 | 0.28035 | 2.932 | 0.02431 * |
| TML - aTPLATE_R | -1.96115 | 0.20411 | -9.608 | < 0.001 *** |
| TML_R - aTPLATE_R | -1.05856 | 0.30951 | -3.420 | 0.00695 ** |
| TML_R - TML | 0.90259 | 0.26948 | 3.349 | 0.00852 ** |

---

Signif. codes: 0 ‘***’ 0.001 ‘**’ 0.01 ‘*’ 0.05 ‘.’ 0.1 ‘ ’ 1

(Adjusted p values reported -- single-step method)

**Figure 4Bi**

Std. Error = Standard Error, aMAP65_R_Rap = MAP-65-1-mCherry-FKBP rapamycin treated, aMAP65_R = MAP-65-1-mCherry-FKBP untreated, MAP65_G = MAP-65-1-GFP untreated, MAP65_G_Rap = MAP-65-1-GFP rapamycin treated

| | Estimate | Std. Error | t value | Pr(>\|t\|) |
|---|---|---|---|---|
| aMAP65_R_Rap - aMAP65_R | 9.16000 | 1.02259 | 8.958 | < 1e-04 *** |
| MAP65_G - aMAP65_R | 0.02429 | 0.16430 | 0.148 | 0.998610 |
| MAP65_G_Rap - aMAP65_R | 1.60071 | 0.35613 | 4.495 | 0.000251 *** |
| MAP65_G - aMAP65_R_Rap | -9.13571 | 1.02291 | -8.931 | < 1e-04 *** |
| MAP65_G_Rap - aMAP65_R_Rap | -7.55929 | 1.07060 | -7.061 | < 1e-04 *** |
| MAP65_G_Rap - MAP65_G | 1.57643 | 0.35707 | 4.415 | 0.000263 *** |

---

Signif. codes: 0 ‘***’ 0.001 ‘**’ 0.01 ‘*’ 0.05 ‘.’ 0.1 ‘ ’ 1

(Adjusted p values reported -- single-step method)

**Figure 5C**

Std. Error = Standard Error, LOL_TPL_TASH3 = LOLITA-GFP in the presence of TPLATE-mCherry-FKBP, MITO-FRB* and TASH3-TagBFP2-FRB* rapamycin treated, LOL_TPL_MITO = LOLITA-GFP in the presence of TPLATE-mCherry-FKBP, MITO-TagBFP2-FRB* and absence of TASH3-TagBFP2-FRB* rapamycin treated

| | Estimate | Std. Error | t value | Pr(>\|t\|) |
|---|---|---|---|---|
| LOL_TPL_TASH3 - LOL_TPL_MITO | 4.0452 | 0.6562 | 6.165 | 1.21e-07 *** |

---

Signif. codes: 0 ‘***’ 0.001 ‘**’ 0.01 ‘*’ 0.05 ‘.’ 0.1 ‘ ’ 1

(Adjusted p values reported -- single-step method)

**Supplemental Figure 3F**

Std. Error = Standard Error, A_Untr = untreated sample, B_Rap_7h = sample treated with rapamycin for 5-7h, C_Asco_6_7_h = sample treated with rapamycin and ascomycin for 6-7h, D_Rap_25_26_h = sample treated with rapamycin for 25-26h, E_Asco_20_26_h = sample treated with rapamycin and ascomycin for 20_26h

| | Estimate | Std. Error | t value | Pr(>\|t\|) |
|---|---|---|---|---|
| B_Rap_7h - A_Untr | 2.2764 | 0.2990 | 7.615 | < 1e-04 *** |
| C_Asco_6_7_h - A_Untr | 5.0048 | 0.5912 | 8.465 | < 1e-04 *** |
| D_Rap_25_26_h - A_Untr | 3.0771 | 0.5294 | 5.813 | < 1e-04 *** |
| E_Asco_20_26_h - A_Untr | 0.4159 | 0.3110 | 1.337 | 0.644720 |
| C_Asco_6_7_h - B_Rap_7h | 2.7284 | 0.6336 | 4.306 | 0.000291 *** |
| D_Rap_25_26_h - B_Rap_7h | 0.8007 | 0.5763 | 1.389 | 0.610981 |
| E_Asco_20_26_h - B_Rap_7h | -1.8605 | 0.3855 | -4.826 | < 1e-04 *** |
| D_Rap_25_26_h - C_Asco_6_7_h | -1.9277 | 0.7696 | -2.505 | 0.086623 . |
| E_Asco_20_26_h - C_Asco_6_7_h | -4.5889 | 0.6394 | -7.177 | < 1e-04 *** |
| E_Asco_20_26_h - D_Rap_25_26_h | -2.6612 | 0.5827 | -4.567 | 0.000108 *** |

---

Signif. codes:  0 '***' 0.001 '**' 0.01 '*' 0.05 '.' 0.1 ' ' 1

(Adjusted p values reported -- single-step method)

p-values multicompView package transformation:

```
multcompLetters
  B     C     D     E     A
 "a"   "b"  "ab"   "c"   "c"
```
